# Proteomic characterization of pilot scale hot-water extracts from the industrial carrageenan red seaweed *Eucheuma denticulatum*

**DOI:** 10.1101/2020.12.14.422673

**Authors:** Simon Gregersen, Margarita Pertseva, Paolo Marcatili, Susan Løvstad Holdt, Charlotte Jacobsen, Pedro J. García-Moreno, Egon Bech Hansen, Michael Toft Overgaard

## Abstract

Seaweeds have a long history as a resource for polysaccharides/hydrocolloids extraction for use in the food industry due to their functionality as stabilizing agents. In addition to the carbohydrate content, seaweeds also contains a significant amount of protein, which may find application in food and feed. Here, we present a novel combination of transcriptomics, proteomics, and bioinformatics to determine the protein composition in two pilot-scale extracts from *Eucheuma denticilatum* (Spinosum) obtained via hot-water extraction. The extracts were characterized by qualitative and quantitative proteomics using LC-MS/MS and a *de-novo* transcriptome assembly for construction of a novel proteome. Using label-free, relative quantification, we were able to identify the most abundant proteins in the extracts and determined that the majority of quantified protein in the extracts (>75%) is constituted by merely three previously uncharacterized proteins. Putative subcellular localization for the quantified proteins was determined by bioinformatic prediction, and by correlating with the expected copy number from the transcriptome analysis, we determined that the extracts were highly enriched in extracellular proteins. This implies that the method predominantly extracts extracellular proteins, and thus appear ineffective for cellular disruption and subsequent release of intracellular proteins. Ultimately, this study highlight the power of quantitative proteomics as a novel tool for characterization of alternative protein sources intended for use in foods. Additionally, the study showcases the potential of proteomics for evaluation of protein extraction methods and as powerful tool in the development of an efficient extraction process.

## 1. Introduction

Seaweeds are known to contain numerous compounds of interest, such as polysaccharides, proteins and other compounds with health beneficial properties such as anti-inflammatory, anti-oxidant, and anti-cancer (Holdt & Kraan, 2011; Leandro et al., 2020). The industry to produce hydrocolloids from seaweed is well established, and the hydrocolloids are used as e.g. stabilizing agents in toothpaste, canned whipped cream, and as meat glue. The production of red carrageenan accounts for 54,000 ton/year and constitutes the majority of the total hydrocolloids sold worldwide (also incl. alginate and agar). Carrageenan is extracted from 212,000 ton dried seaweed, and brings in a value of 530 million USD (Porse, 2018). *Eucheuma denticulatum* is among the most cultivated and harvested red seaweed species for the carrageenan industry. However, at present carrageenan is extracted in a process, which extracts carrageenan as the only compound whereas proteins and other compounds are not extracted. The most common industrial method to extract carrageenan from *Eucheuma denticulatum* uses hot water at high pH. If further extraction of other compounds such as proteins could be made prior to or as part of the industrial hot water extraction without compromising the existing carrageenan extraction, this could be of interest, since the amount of biomass available is large. Proteins from *E. denticulatum* were shown to constitute only 3.8% of dry biomass, but were of high quality with respect to their amino acid profile (Naseri, Jacobsen, et al., 2020). Moreover, the obtained proteins are comparable to beef in regard to the branched chained amino acids (i.e. leucine, isoleucine, and valine) that are of interest due to their muscle building properties.

In addition to the general health benefits from ingestion (Gomez-Zavaglia, Prieto Lage, Jimenez-Lopez, Mejuto, & Simal-Gandara, 2019; Peñalver et al., 2020), seaweed may also be a source of bioactive peptides that could exhibit a direct biological purpose or be utilized as functional food ingredients. These peptides can be released through bio-processing of proteins extracts using e.g. enzymatic hydrolysis or fermentation (Admassu, Gasmalla, Yang, & Zhao, 2018). In the past decade, peptides derived from seaweed proteins with e.g. renin-inhibitory (Fitzgerald et al., 2012), ACE-inhibitory (Furuta, Miyabe, Yasui, Kinoshita, & Kishimura, 2016), antioxidant (Cian, Garzón, Ancona, Guerrero, & Drago, 2015), and antidiabetic (Harnedy & FitzGerald, 2013b) activities have been identified. Common for all bioactive peptides is that they were identified in enzymatic hydrolysates by a non-targeted trial-and-error approach. This methodology, commonly employed in the food industry, requires numerous costly and time-demanding steps of hydrolysis, separation, isolation, identification, and finally *in vitro* or *in vivo* verification of activity. In contrast, an orthogonal approach utilizing bioinformatic prediction of bioactive peptides, is gathering increased attention (Tu, Cheng, Lu, & Du, 2018). This method reduces cost and work load tremendously, and allows for targeted peptide release by enzymatic hydrolysis. With recent advances in bioinformatic prediction of peptide functionality (García-Moreno, Jacobsen, et al., 2020; Mooney, Haslam, Holton, Pollastri, & Shields, 2013; Mooney, Haslam, Pollastri, & Shields, 2012; Olsen et al., 2020; Panyayai et al., 2019), and the growing availability of peptide databases (Chen et al., 2013; Liu, Baggerman, Schoofs, & Wets, 2008; Minkiewicz, Iwaniak, & Darewicz, 2019; G. Wang, Li, & Wang, 2009), the primary prerequisite for the analysis is the availability of protein sequences and quantitative information on protein composition. Recently, we employed quantitative proteomics for identification of abundant proteins followed by bioinformatic prediction (EmulsiPred source code freely available at https://github.com/MarcatiliLab/EmulsiPred) to identify a number of highly functional emulsifier peptides from potato (García-Moreno, Gregersen, et al., 2020) as well predicting probable emulsifier and antioxidant peptides in hydrolysates from fish processing side streams following LC-MS/MS analysis (Jafarpour, Gomes, et al., 2020; Jafarpour, Gregersen, et al., 2020). Nevertheless, proteomic quantification of the starting material is an absolute necessity in order to maximize the yield of peptide release. Here, we present a proteomic characterization of two industrially relevant, pilot-scale extracts from *E. denticulatum* obtained by hot-water extraction. Protein identification is based on a de novo transcriptome assembly for creating a novel reference proteome. Furthermore, we present a novel approach for quantifying proteins based on non-tryptic peptides, and correlate protein abundance with quantitative transcriptomics. Using bioinformatic prediction of protein subcellular origin, we are able to determine enrichment of certain protein classes in the extracts.

## 2. Materials and Methods

### 2.1. Materials

Two *Eucheuma denticulatum* protein extracts obtained using near-neutral, hot-water extraction were supplied by the global food ingredient provider CP Kelco. Protein extract A was obtained by dispersing the raw seaweed in deionized water (pH adjusted to 8.9 with sodium carbonate) and applying continuous stirring at 95°C for 5 h. The slurry was subsequently filtered in a Büchner funnel followed by diafiltration using a 300 kDa MWCO membrane. The retentate was washed with three volumes of 0.9% sodium chloride in deionized water, and all permeates were subsequently pooled. The pooled permeate was then concentrated using a 1 kDa MWCO membrane, and the retentate lyophilized to yield the final protein extract A. Protein extract B was obtained similarly to extract A, but with stirring at 90°C for 16 h before filtering, diafiltration, concentration, and lyophilization. Furthermore, the lyophilized retentate was dissolved in deionized water, the pH was adjusted to 2.9 with nitric acid, and the mixture was stirred at room temperature for 1 h. Precipitated protein was isolated by centrifugation and washed twice with isopropanol before air drying and lyophilization to yield the final protein extract B. The total protein content of protein extracts A and B (by Kjeldahl-N) was 7.1% and 70% (w/w), respectively, using a nitrogen-to-protein conversion factor of 6.25 (CP Kelco supplied information). All chemicals used were of analytical grade.

### 2.2. Total soluble protein

Protein extracts A and B were solubilized to an estimated protein concentration of 2 mg/mL in ddH_2_O and in 200 mM NH_4_HCO_3_ with 0.2% SDS for maximal solubilization compatible with the Qubit protein assay. Following solvent addition, samples were vortexed for 30 s, sonicated for 30 min, and left overnight on a Stuart SRT6 roller mixer (Cole-Parmer, UK). The next day, samples were sonicated for 30 min, left on a roller mixer for 60 min, and centrifuged at 3,095 RCF (ambient temperature) for 10 min in a 5810 R centrifuge (Eppendorf, Germany), prior to aliquoting the supernatant. The total soluble protein content of the samples in both solvents, was quantified using Qubit protein assay (Thermo Scientific, Germany) according to the manufacturer guidelines.

### 2.3. 1D-SDS-PAGE and in-gel digestion

Protein extracts A and B were solubilized with 2% SDS in 200 mM ammonium bicarbonate (pH 9.5) to a final protein/peptide concentration of 2 mg/mL based on protein content by Kjeldahl-N. Alkaline buffer with detergent was used to maximize protein solubilization. Solubilization was further promoted by. Samples were vortexed for 2 min, sonicated for 30 min, and subsequently centrifuged at 3,095 RCF for 15 min to precipitate solids. SDS-PAGE analysis was performed on precast 4-20% gradient gels (GenScript, USA) in a Tris-MOPS buffered system under reducing conditions according to manufacturer guidelines. Briefly, 20 μg protein/peptide was mixed with reducing (final DTT concentration 50 mM) SDS-PAGE sample buffer and subsequently denatured at 95 °C for 5 min prior to loading on the gel. As molecular weight marker, PIERCE Unstained Protein MW Marker P/N 26610 (ThermoFisher Scientific, USA) was used. Protein visualization was achieved by using Coomassie Brilliant Blue G250 staining (Sigma-Aldrich, Germany) and imaging with a ChemDoc MP Imaging System (Bio-Rad, USA).

Proteins were in-gel digested according to Shevchenko et al. (Shevchenko, Wilm, Vorm, & Mann, 1982) and Fernandez-Patron et al. (Fernandez-Patron et al., 1995), as previously described (García-Moreno, Gregersen, et al., 2020). Briefly, each gel lane from the gradient gel was excised with a scalpel and divided into 6 fractions guided by the MW marker (<14kDa; 14-25kDa; 25-45kDa; 45-66kDa; 66-116kDa; >116kDa). Individual fractions were cut into 1×1 mm pieces before being subjected to washing, reduction with DTT, Cys alkylation with iodoacetamide, and digestion with sequencing grade modified trypsin (Promega, Madison, WI, USA). Following digestion, peptides were extracted, dried down by SpeedVac, and suspended in 0.1% (v/v) formic acid (FA), 2% acetonitrile (ACN) (v/v). Next, peptides were desalted using StageTips (Fernandez-Patron et al., 1995; Rappsilber, Mann, & Ishihama, 2007), dried down by SpeedVac, and finally suspended in 0.1% (v/v) FA, 2% ACN (v/v) for LC-MS/MS analysis.

### 2.4. De novo transcriptome assembly

The transcriptome of *E. denticulatum* was downloaded from the NCBI SRA database (https://www.ncbi.nlm.nih.gov/sra/SRX2653634). The raw reads were preprocessed by Trimmomatic software to filter short sequences (less than 36 bp) and to trim low-quality ends (Bolger, Lohse, & Usadel, 2014). Processed reads were then assembled *de novo* into contigs using Trinity with default parameters (Grabherr et al., 2011). Overall, 9458 contigs were assembled with an average length of 1021 bp.

### 2.5. Transcript annotation, abundance estimation and protein database construction

The potential protein-coding sequences were predicted by TransDecoder based on the length of open reading frames and nucleotide composition (Grabherr et al., 2011). Candidate sequences were annotated by BlastP and BlastX search against SwissProt database (Madden, 2013) with the cutoff E-value of 1E-5 as well as by HMMER (Finn, Clements, & Eddy, 2011) search against Pfam database (El-Gebali et al., 2018; Finn et al., 2010). An alignment E-value of 1E-5 means that a homology hit has a 1 in 100,000 probability of occurring by chance alone, therefore we chose this threshold to get only high-quality homologous proteins hits.

The abundance of the transcripts (transcripts per megabase, TPM) was calculated by re-aligning reads to the assembled contigs using RSEM (RNA-Seq by Expectation-Maximization) estimation method included in Trinity software (Grabherr et al., 2011). Obtained transcript abundance matrix was joined with Blastp-annotated transcripts to attain a list of highly expressed proteins.

### 2.6. Prediction of subcellular localization using deepLoc

All proteins in the final database were analyzed by deepLoc (Almagro Armenteros, Sønderby, Sønderby, Nielsen, & Winther, 2017) using the freely available web-tool (http://www.cbs.dtu.dk/services/DeepLoc/index.php). All searches were performed using the BLOSUM62 protein encoding to achieve a probability based subcellular localization for use in enrichment analysis on both transcriptome and protein level.

### 2.7. LC-MS/MS analysis

Tryptic peptides were analyzed by an automated LC–ESI–MS/MS consisting of an EASY-nLC system (Thermo Scientific, Bremen, Germany) on-line coupled to a Q Exactive mass spectrometer (Thermo Scientific) via a Nanospray Flex ion source (Thermo Scientific), as previously reported (García-Moreno, Gregersen, et al., 2020). Separation of peptides was achieved by use of an Acclaim Pepmap RSLC analytical column (C18, 100 Å, 75 μm. × 50 cm (Thermo Scientific)). Instrumental settings, solvents, flows, gradient, and acquisition method were identical to what was described previously.

### 2.8. Proteomics data analysis

Protein identification and quantification was performed using MaxQuant 1.6.0.16. (Cox & Mann, 2008; Tyanova, Temu, Sinitcyn, et al., 2016) using the de-novo proteome assembled from the transcriptomic analysis. Initially, standard settings were employed using specific digestion (Trypsin/P, 2 missed cleavages allowed, minimum length 7 AAs) and false discovery rate (FDR) of 1% on both peptide and protein level. FDR was controlled using reverse decoy sequences and common contaminants were included. Protein quantification was obtained with including both unique and razor peptides. Samples were analyzed as six fractions with boosted identification rates by matching between runs and dependent peptides enabled. The iBAQ algorithm (Schwanhüusser et al., 2011) was used for relative in-sample protein quantification. iBAQ intensities were normalized to the sum of all iBAQ intensities after removal of reverse hits and contaminants, to obtain the relative iBAQ (riBAQ), as previously described (García-Moreno, Gregersen, et al., 2020; Shin et al., 2013).

MS-data were furthermore analyzed both semi-specifically (tryptic *N*- or *C*-terminus) and unspecifically (no terminal restrictions) in MaxQuant. All settings were maintained except for applying unspecific digestion with peptide length restrictions from 4 to 65 AAs. Additional unspecific searches with peptide and protein level FRD of 5% and 10% as well as semi-specific searches with peptide and protein level FRD of 5% was conducted to increase identification rates and sequence coverage for comparison and data quality assessment.

Relative quantification with iBAQ employs strict tryptic restrictions to peptide termini and consequently, this type of quantifications is not possible for semi-specific and unspecific searches. In order to compare and evaluate the semi-specific and unspecific results, we introduced two additional quasi-quantitative relative metrics: i) relative intensity, I_rel_ and ii) length-normalized relative intensity, I^L^_rel_. They were defined as:

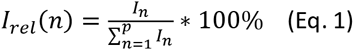

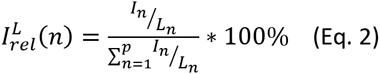

Where I_n_ is the intensity of protein n of p quantified proteins in a given sample and L_n_ is the length of protein n, based on the processed protein database. For evaluation of the two metrics, relative protein abundance was plotted as scatter plots between the different analysis conditions and the Pearson correlation coefficient (PCC) was calculated in Perseus (Tyanova, Temu, Sinitcyn, et al., 2016).

For final protein quantification, MS data were analyzed as both tryptic and semi-tryptic digests using the following optimized search criteria: Peptides per protein ≥ 2 (razor and unique), protein FDR = 0.05, unmodified peptide score > 40, peptide FDR = 0.005. Match between runs and dependent peptides were both disabled. This was done to alleviate false positive identifications and increase quantitative validity. Increasing FDR to 5% for the tryptic analysis did not affect identification and quantification due to the applied score threshold.

### 2.9. Comparative analysis of transcriptomic and proteomic data

Comparative analysis was done on both the protein and subcellular levels. To estimate molar transcript abundance, we calculated the relative TPM (rTPM) for the individual proteins to the sum of TPMs for all 1628 proteins in the database. Using the predicted subcellular localization, we then estimated the relative distribution of proteins based on the transcriptome using rTPM. Finally, we correlated the transcriptome-based protein distribution with the actual protein distribution for the extracts in a relative, quantitative manner.

### 2.10. Data analysis and visualization

Statistical and correlation analysis of transcriptome and MS data was performed in Perseus 1.6.1.3 (Tyanova & Cox, 2018; Tyanova, Temu, Sinitcyn, et al., 2016). Venn diagrams were plotted with jvenn (Bardou, Mariette, Escudié, Djemiel, & Klopp, 2014). Additional data visualization was obtained using OriginPro 8.5.0 SR1 (OriginLab Corporation, Northampton, MA, USA) and figures assembled in their final form using INKSCAPE version 0.92.3 (https://inkscape.org/).

## 3. Results and Discussion

### 3.1. Transcriptome assembly, protein annotation, and subcellular localization

The transcriptome of *Eucheuma denticulatum* was de novo assembled using publicly deposited transcriptome data at NCBI SRA database. The quality of the assembly was estimated by basic contig statistics and percentage of the remapped reads. Both metrics indicated a high quality of the assembly with an N50 value of 1891bp (Table A.1) and more than 90% of the reads mapped back to the contigs (Table A.2). Based on the transcriptomic information, an *E. denticulatum* protein database was constructed for subsequent mass-spectrometry (MS) analysis. First, the protein-coding sequences were predicted and identified their by BlastX and BlastP search as well as their protein family by searching against Pfam database. Then the transcript expression level was calculated in terms of transcripts per kilo megabase (TPM) and removed proteins with TPM below 100, which resulted in 1628 proteins retained for the database. The TPM threshold was applied in order to filter out any potentially erroneous reads. Although this may in fact also filter some proteins with low copy numbers from the database, the primary objective was to identify highly expressed and extracted proteins, and consequently do not regard this to have substantial influence. A full list of protein accessions and their associated TPMs, rTPMs, Pfam functions, BlastX targets, and BlastP targets can be found in Table A.3 and in the linked Mendeley data repository. The de-novo protein database for *E. denticulatum* can be found in .fasta format in Table A.4 as well as in the Mendeley data repository.

Although homology-inferred annotation using BLAST can indicate potential functions and localizations for the individual proteins, extraction of potential functions and subcellular localization on the proteome level is a tedious task. Additionally, as only verified Uniprot/Swiss-Prot proteins were included, the resulting annotations were of suboptimal quality (Table A.3) due to the lack of verified annotations on related and comparable species to *E. denticulatum*. Consequently, a bioinformatic prediction of subcellular localization on the individual protein level was used. This data type is easily binnable for large proteomes. As the DeepLoc neural network was developed for eukaryotic proteins with little or no available homology data (Almagro Armenteros et al., 2017), this directly applies to the case of this study. For the entire proteome, DeepLoc achieved a localization probability of 0.63 ± 0.21 (Figure A.1).

### 3.2. 1D SDS-PAGE analysis and protein quality assessment

Both protein extracts display absence of distinct proteins bands and an apparent smear along the gel concentrating in the low MW range, as seen from 1D SDS-PAGE analysis in Figure 1. This is in contrast to previous studies on *E. denticulatum* protein extracts (Rosni et al., 2015), where distinct protein bands were observed and the low MW concentrated smear was absent. The significant difference in protein appearance by SDS-PAGE may be directly ascribed to the extraction method, as the authors here used a more elaborate protocol including organic (phenol) solvents as well as reducing conditions. Their approach may be significantly better for efficient extraction of intact proteins from the whole seaweed, but is not feasible on an industrial scale.

**Figure 1:**
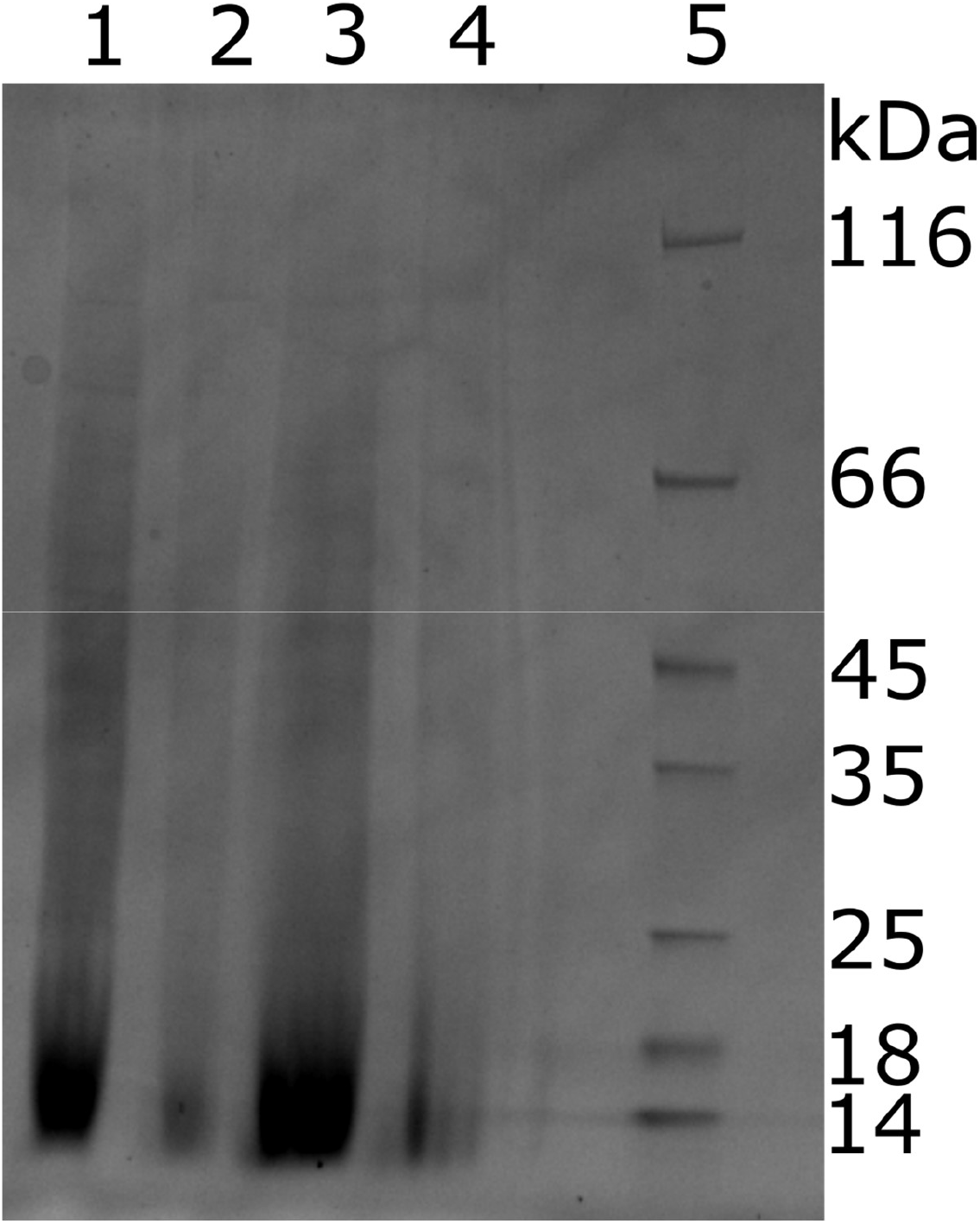
SDS-PAGE of *E. denticulatum* protein extracts investigated in this study. Protein loading is based on supplied protein content of 7.1% and 70% for extract A and B, respectively. 1: Extract A, 100 μg. 2: Extract A, 20 μg. 3: Extract B, 100 μg. 4: Extract B, 20 μg. 5: MW Marker.

The overall appearance of both extracts analyzed, however, are quite similar. The lack of distinct protein bands could potentially indicate partial hydrolysis during extraction using high temperature under alkaline conditions, as employed for both extraction methods. In addition, the extraction methodology employed may also result in co-extraction of other cellular moieties, which could interfere with electrophoresis and ultimately resulting in the observed smears. This has been reported for co-extracted lipids (Simões-Barbosa, Santana, & Teixeira, 2000; W. Wang et al., 2004), carbohydrates (Chart & Rowe, 1991; Hashimoto & Pickard, 1984), and DNA (Park, Kim, Choi, Grab, & Dumler, 2004). Further modification of proteins (e.g. glycoproteins) may also add to the smearing observed on SDS-PAGE (Elliott et al., 2004; Møller & Poulsen, 2009; Sparbier, Koch, Kessler, Wenzel, & Kostrzewa, 2005).

In order to estimate the accuracy of the total protein by Kjeldahl-N analysis, we determined the soluble protein content in both aqueous solution and a slightly alkaline buffer with added detergent using Qubit protein assay (Table 1). From here, it is evident that the Kjeldahl-based total protein in fact correlates quite well with the soluble protein content – at least when solubilized in an alkaline buffer with detergent. A nitrogen-to-protein conversion factor of 6.25, the “Jones factor”, is commonly employed in food protein science and has been so for 90 years (Jones, 1931; Salo-Väänänen & Koivistoinen, 1996). Nevertheless, the universal conversion factor has been subject to several investigations, and species-dependent conversion factors are commonly recommended (Mariotti, Tomé, & Mirand, 2008). For seaweeds in particular, the factor can still vary significantly, but as no factor is available for *E. denticulatum*, a general conversion factor of 5.0 can be applied (Angell, Mata, de Nys, & Paul, 2016). By doing so, and thereby lowering the protein content by 20% (Table 1), the Kjeldahl-N method now underestimates the protein content compared to Qubit – in particular for extract B. In this respect, it is worth considering that the conversion factor is representative of the total organism proteome. Additionally, the non-protein nitrogen content of the extract is undetermined, and may also influence both the Kjeldahl-N and the Qubit outputs to some degree.

**Table 1:**
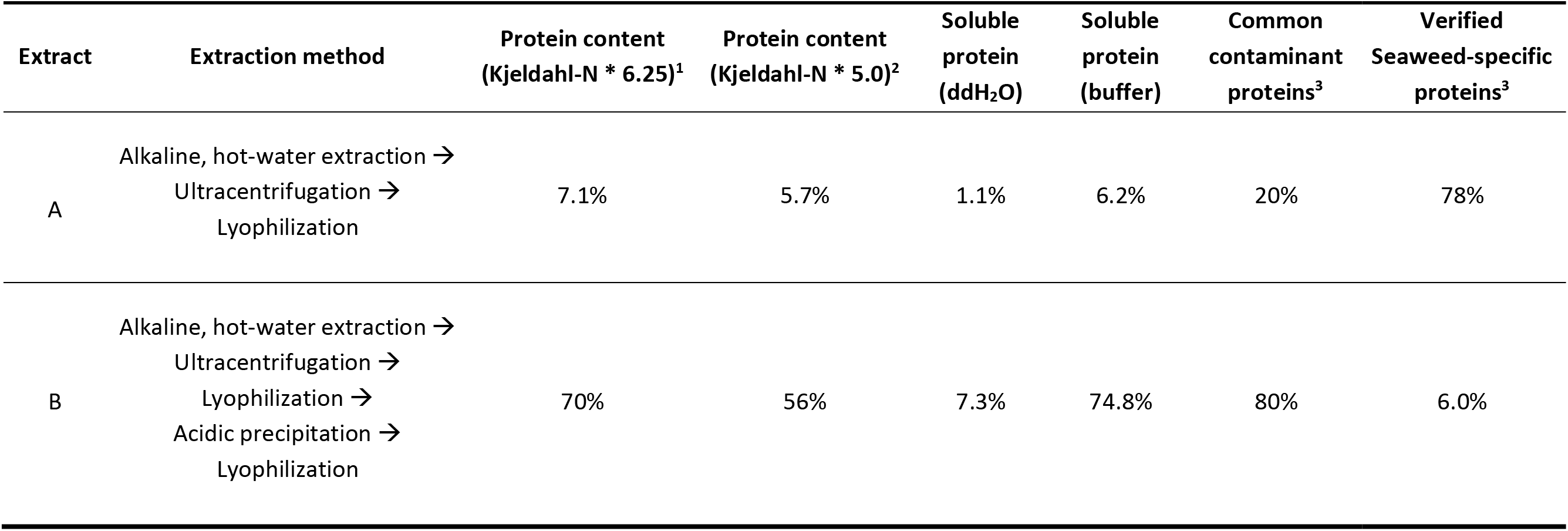
General characteristics for the two *E. denticulatum* extracts analyzed in this work including total protein and soluble protein content. ^1^Total protein by Kjeldahl-N was supplied by CP Kelco. ^2^Calculated based on supplied protein content (^1^) using a conversion factor of 5.0 (Angell et al., 2016). ^3^Sum of relative abundance for common contaminant proteins and verified seaweed specific proteins identified in MaxQuant by I^L^_rel_ for semi-specific analysis with optimized parameters, prior to any filtering, but after removing trypsin (Stage 1).

It is also evident that the aqueous solubility of the protein in the extracts is quite low (11-15% of the total protein), whereas a slightly alkaline buffer with a low amount of detergent practically fully solubilizes the protein (6-fold and 10-fold solubility increase for extract A and B, respectively). This also correlates well with the physical appearance of the solubilized extracts following centrifugation (Figure A.2), where a significantly higher amount of solid precipitate is visible in the aqueous solutions. Nevertheless, smear and apparent lack of intact high MW protein from SDS-PAGE must be taken into consideration for protein quantification and in the evaluation of the protein extracts as source for further processing as well as potential release of bioactive peptides.

### 3.3. Identification and quantification of peptides and proteins by LC-MS/MS

Initially, we applied an iterative process where different *in silico* digestion methods (i.e. specific, semi-specific, and unspecific digestion), peptide- and protein-level FDR, and number of identified peptides per protein were attempted. This was done not only to identify the optimal parameters for analysis, but also to investigate the feasibility of applying the two specified quantitative metrics. The iterative process was of utmost importance, as the sample quality and especially the number of identified peptides and proteins for the extracts was low. A low number of peptide identifications significantly affects protein identification and quantification via the impact on FRD-controlled thresholds. This is ultimately an inherent property of the peptide scoring algorithm. MaxQuant employs the Andromeda search engine, in which peptide score is not only based on PEP, but also on the intensity of a given feature (Cox et al., 2011; Tiwary et al., 2019; Tyanova, Temu, & Cox, 2016). Consequently, high intensity features with significant PEP (i.e. potential false positives), which in other studies may have been filtered out, will obtain a sufficiently high peptide score and be used in protein quantification. Ultimately, this leads to false identification of proteins with a significant relative abundance, which impairs further analysis. By applying more stringent thresholds on both peptide and protein level, this is alleviated to some extent. Nevertheless, it may be needed to inspect and evaluate PEPs rather than apply threshold filtering on peptide score alone, as PEP relies solely on PSM and sequence-dependent features. This aspect is thoroughly discussed and evaluated in Appendix A.

By applying the optimized search parameters, a total of 66 proteins across both extracts and analysis methods (tryptic and semi-specific) following filtering of trypsin and reverse hits (Stage 1) were identified and quantified (Table 2). Extract B is highly contaminated since 80% (based on I^L^_rel_ for semi-tryptic analysis) of all identified proteins were constituted by common contaminants (Table 1), primarily keratins. On the other hand, extract A “only” contained 20%. Although common contaminants are usually filtered out prior to quantification, the magnitude is noteworthy. In total, merely 40 proteins were identified across both extracts and analysis conditions, following filtering of common contaminants and subsequent re-quantification (Stage 2, Tables A.5; A.6). Semi-specific analysis resulted in identification of four additional proteins (one in extract A and three in extract B), whereof one (c1275_g1_i1.p1) constitutes more than half of the Stage 2 protein by I^L^_rel_ in extract B. Furthermore, 11 proteins were not identified by this approach (four in extract A, three in extract B and four identified in both extracts using tryptic conditions), but none of these were of high abundance. From plotting relative abundance by both riBAQ and I^L^_rel_ (Figure A.3), a correlation was seen within each extract (PPC = 0.99-1.0 for extract A; PPC = 0.19-0.95 for extract B), but the semi-specific analysis of extract B correlated poorly with the tryptic analysis. The correlation between extracts was even worse (PPC = 0.14-0.55), indicating that the stringent quality parameters applied for automatic filtering, were not fully capable of cleaning the data from bad peptide spectrum matches (PSMs) and dubious protein identifications.

**Table 2:** Relative protein abundance of *E. denticulatum* extracts A and B (after filtering of trypsin) following initial quantification (Stage 1) with optimized search parameters by I^L^_rel_ and riBAQ for both tryptic and semi-specific analysis. Proteins are divided in common contaminants (Stage 1 filtered proteins), false positive identifications/contaminants (Stage 2 filtered protein), and final, verified proteins (Stage 3). Common contaminants are annotated using their UniProt accession number. NQ: Protein not quantified in the specific sample using the specific analysis method.

| Stage 1 |  |  |  |  |  |  |  | Stage 2 |  |  |  |  |  |  | Stage 3 |  |  |  |  |  |  |
| --- | --- | --- | --- | --- | --- | --- | --- | --- | --- | --- | --- | --- | --- | --- | --- | --- | --- | --- | --- | --- | --- |
| Protein IDs | I<sub>rel</sub> A<br>tryp | riBAQ<br>A | I<sub>rel</sub> A<br>semi | I<sub>rel</sub> B<br>tryp | riBAQ<br>B | I<sub>rel</sub> B<br>semi | Contaminant<br>ID | Protein IDs | I<sub>rel</sub> A<br>tryp | riBAQ<br>A | I<sub>rel</sub> A<br>semi | I<sub>rel</sub> B<br>tryp | riBAQ<br>B | I<sub>rel</sub> B<br>semi | Protein IDs | I<sub>rel</sub> A<br>tryp | riBAQ<br>A | I<sub>rel</sub> A<br>semi | I<sub>rel</sub> B<br>tryp | riBAQ<br>B | I<sub>rel</sub> B<br>semi |
| CON_ENSEMBL | NQ | NQ | NQ | 0.1% | 0.1% | NQ | Keratin | rf1c10492_g1_i1.p1 | NQ | NQ | NQ | 0.1% | 0.1% | NQ | rf1c1505_g2_i1.p1 | 8.6% | 11.8% | 13.4% | 1.2% | 1.5% | 1.7% |
| CON_O43790 | NQ | NQ | NQ | 0.7% | 0.5% | 0.6% | Keratin | rf1c1275_g1_i1.p1 | NQ | NQ | NQ | NQ | NQ | 11.1% | rf1c1613_g1_i1.p1 | NQ | NQ | 0.0% | 0.0% | 0.0% | 0.0% |
| CON_P02533 | 0.1% | 0.1% | 0.0% | 1.9% | 1.5% | 1.8% | Keratin | rf1c1294_g1_i1.p1 | 0.0% | 0.0% | NQ | 0.0% | 0.0% | NQ | rf1c17304_g1_i1.p1 | 2.6% | 2.3% | 2.5% | 0.3% | 0.2% | 0.2% |
| CON_P02662 | NQ | NQ | NQ | 0.1% | 0.1% | NQ | α-S1-casein | rf1c1357_g1_i1.p1 | NQ | NQ | NQ | 0.0% | 0.0% | NQ | rf1c17615_g1_i1.p1 | 0.1% | 0.1% | NQ | NQ | NQ | NQ |
| CON_P02666 | 0.0% | 0.0% | 0.0% | 2.2% | 3.8% | 1.8% | β-casein | rf1c17161_g1_i1.p1 | NQ | NQ | NQ | 0.1% | 0.0% | NQ | rf1c231_g1_i1.p1 | 0.1% | 0.1% | NQ | NQ | NQ | NQ |
| CON_P02754 | 6.5% | 4.7% | 6.9% | 5.2% | 3.7% | 4.9% | β-lactoglobulin | rf1c17201_g1_i1.p1 | 0.0% | 0.0% | NQ | NQ | NQ | NQ | rf1c2364_g1_i1.p1 | 0.1% | 0.1% | 0.1% | 0.5% | 0.4% | 0.4% |
| CON_P02768 | 0.0% | 0.0% | 0.0% | 0.0% | 0.0% | 0.0% | Albumin | rf1c17231_g1_i1.p1 | 0.0% | 0.0% | 0.0% | 0.3% | 0.3% | 0.2% | rf1c2556_g1_i1.p1 | 0.0% | 0.0% | NQ | 0.0% | 0.0% | NQ |
| CON_P02769 | 0.0% | 0.0% | 0.0% | 1.5% | 1.1% | 1.3% | Albumin | rf1c2788_g1_i1.p1 | NQ | NQ | NQ | NQ | NQ | 0.2% | rf1c3760_g1_i1.p1 | NQ | NQ | 0.7% | NQ | NQ | NQ |
| CON_P04264 | 5.5% | 5.6% | 4.6% | 33.3% | 33.1% | 26.7% | Keratin | rf1c3249_g1_i1.p1 | 0.0% | 0.0% | NQ | NQ | NQ | NQ | rf1c4090_g1_i1.p1 | 0.1% | 0.1% | 0.1% | 0.0% | 0.0% | NQ |
| CON_P08779 | 0.0% | 0.0% | 0.0% | 0.3% | 0.2% | 0.2% | Keratin | rf1c4757_g1_i1.p1 | NQ | NQ | NQ | 0.1% | 0.1% | 0.2% | rf1c4354_g1_i1.p1 | 3.3% | 4.1% | 3.1% | 0.0% | 0.0% | 0.0% |
| CON_P13645 | 6.2% | 6.9% | 5.5% | 19.3% | 21.1% | 16.8% | Keratin | rf1c4921_g1_i1.p1 | NQ | NQ | NQ | 0.0% | 0.0% | 0.1% | rf1c4671_g1_i1.p2 | 0.3% | 0.4% | 0.2% | NQ | NQ | NQ |
| CON_P13647 | 0.2% | 0.2% | 0.2% | 1.8% | 1.6% | 0.9% | Keratin | rf1c5168_g1_i1.p1 | NQ | NQ | NQ | 0.0% | 0.0% | 0.1% | rf1c5232_g1_i1.p1 | 0.9% | 0.9% | 0.8% | NQ | NQ | NQ |
| CON_P19013 | NQ | NQ | 0.0% | 0.2% | 0.1% | 0.1% | Keratin | rf1c5952_g1_i1.p1 | NQ | NQ | NQ | NQ | NQ | 0.1% | rf1c6313_g1_i1.p1 | 27.2% | 25.3% | 25.2% | 0.2% | 0.2% | 1.4% |
| CON_P35527 | 1.1% | 1.3% | 1.1% | 17.2% | 19.7% | 16.2% | Keratin | rf1c6797_g1_i1.p1 | NQ | NQ | NQ | 0.0% | 0.0% | 0.0% | rf1c6373_g1_i1.p1 | 0.0% | 0.0% | 0.0% | 0.0% | 0.0% | NQ |
| CON_P35908 | 2.3% | 1.8% | 1.9% | 6.1% | 4.7% | 5.9% | Keratin | rf1c6825_g1_i1.p3 | 1.4% | 0.9% | 1.3% | 2.8% | 1.7% | 2.1% | rf1c6458_g1_i1.p1 | 1.1% | 1.8% | 1.4% | NQ | NQ | NQ |
| CON_P48668 | 0.0% | 0.0% | 0.0% | 0.6% | 0.6% | 1.3% | Keratin | rf1c6945_g1_i1.p2 | 0.0% | 0.0% | 0.0% | 0.2% | 0.2% | 0.2% | rf1c6656_g1_i1.p1 | 0.0% | 0.0% | 0.0% | 0.3% | 0.3% | 0.3% |
| CON_P78386 | 0.0% | 0.0% | NQ | 0.1% | 0.1% | 0.1% | Keratin | rf1c8389_g1_i1.p1 | 0.0% | 0.0% | NQ | 0.1% | 0.0% | NQ | rf1c6834_g1_i1.p3 | 0.0% | 0.0% | NQ | 0.0% | 0.0% | NQ |
| CON_Q04695 | NQ | NQ | 0.0% | 0.2% | 0.1% | 0.2% | Keratin |  |  |  |  |  |  |  | rf1c6963_g2_i1.p1 | 0.1% | 0.1% | 0.0% | 0.0% | 0.0% | 0.0% |
| CON_Q14525 | 0.0% | 0.0% | 0.0% | 0.1% | 0.0% | 0.0% | Keratin |  |  |  |  |  |  |  | rf1c7052_g1_i1.p1 | 27.0% | 25.1% | 25.8% | 1.6% | 1.5% | 1.5% |
| CON_Q5D862 | 0.0% | 0.0% | NQ | 0.0% | 0.0% | NQ | Filaggrin-2 |  |  |  |  |  |  |  | rf1c7052_g1_i2.p1 | 3.5% | 3.9% | 3.2% | 0.4% | 0.4% | 0.4% |
| CON_Q6KB66 | NQ | NQ | NQ | 0.0% | 0.0% | NQ | Keratin |  |  |  |  |  |  |  | rf1c7216_g1_i1.p1 | 0.3% | 0.3% | 0.4% | NQ | NQ | NQ |
| CON_Q9UE12 | 0.0% | 0.0% | NQ | 0.6% | 0.5% | 0.5% | Keratin |  |  |  |  |  |  |  | rf1c8421_g1_i1.p1 | 1.2% | 2.0% | 1.2% | NQ | NQ | NQ |
| CON_Q9NSB2 | NQ | NQ | NQ | 0.1% | 0.0% | 0.0% | Keratin |  |  |  |  |  |  |  | rf1c926_g1_i1.p1 | 0.0% | 0.0% | 0.0% | NQ | NQ | NQ |
| CON_Q7Z3Y8 | NQ | NQ | NQ | 0.0% | 0.0% | 0.0% | Keratin |  |  |  |  |  |  |  |  |  |  |  |  |  |  |
| CON_Q86YZ3 | NQ | NQ | NQ | 0.0% | 0.0% | 0.0% | Hornerin |  |  |  |  |  |  |  |  |  |  |  |  |  |  |
| CON_Q8IUT8 | NQ | NQ | NQ | NQ | NQ | NQ | Keratin |  |  |  |  |  |  |  |  |  |  |  |  |  |  |

Identified “outliers” (Tables A.5; A.6) that did not correlate between extracts (i.e. are suddenly highly enriched in extract B) may in fact be contaminants with some homology to the *E. deticulatum* proteome (further details are presented in the Appendix A). For instance, in the tryptic analysis of extract B, c6825_g1_i1.p3 is highly abundant but only identified by two peptides, which both map to histones from e.g. humans. Histone was also the BLASTX target (*Xenopus laevis* (African clawed frog) histone H2AX) as well as the predicted function by Pfam (Table A.3). Consequently, and because it was very low abundance in extract A, this was ascribed as contaminant to the extract and not originating from the seaweed. Although histones were bound to be identified in *E. denticulatum*, homologues from other organisms would bias quantification and it was consequently excluded. Furthermore, the highly abundant protein identified by semi-specific analysis of extract B only (c1275_g1_i1.p1), was also identified by only two peptides. As the protein score of 11.8 was very low (see Appendix A and Table A.6), and the posterior error probability (PEP) was significant (PEP>0.05), these were regarded bad PSMs and the protein ID was deemed false positive. Based on these observations, manual inspection and validation was performed in order to apply a final filtering step using the rationale described above. In the filtering, significant weight was put on evaluation of PEP rather than peptide score, as low scoring peptides (< 40) were pre-filtered in the optimized search parameters (see Appendix A for further details). Filtered proteins, along with the rationale for their exclusion, can be found in Table A.7 and proteins are listed under Stage 2 in Table 2.

Following filtering, verified proteins were re-quantified (Stage 3) the list of identified proteins was reduced from 40 to 23 proteins across extracts and conditions (Table 3). The stringent parameters applied in data analysis, as well requirements for inclusion in the final list, fully alleviated the problem of new and significantly abundant proteins showing up in extract B (see Table 2 and Figure A.3), as no proteins exclusive for extract B, were observed (Figure 2B). Nine proteins were observed exclusively in extract A, but this may be explained as loss during the extended processing for extract B. Extended processing may also be a likely explanation for the extract B exclusive peptides identified (Figure 2A). Furthermore, all nine proteins are of somewhat low abundance (I^L^_rel_ < 2%), and do not affect the overall protein distribution significantly. Interestingly, the 9 proteins identified in both extracts using both analyses approaches, constituted > 93% of the verified protein in extract A and > 99% of the verified protein in extract B (by I^L^_rel_). In fact, three proteins (c6313_g1_i1.p1, c7052_g1_i1.p1, and c1505_g2_i1.p1) constitute more than 75% of the total protein identified in both extracts (Table 3). Furthermore, an isoform of c7052_g1_i1 (c7052_g1_i2), which only differs in the C-terminal region of the protein, was also identified in significant abundance. If included, the proteins constitute > 80% of the verified seaweed-specific protein in both extracts. With MW in the range 16-24 kDa, all three (four) proteins correlated well with the observations from SDS-PAGE (Figure 1), even though no clear protein bands were observed. This indicated that these three (four) proteins in particular may be of certain interest as potential sources of e.g. bioactive peptides. Protein sequences and experimental sequence coverage for the three major proteins are shown in Figure 3. From BlastP against verified proteins in UniProtKB/Swiss-Prot (Table S3), c7052_g1_i1.p1 (as well as the isoform) shows some homology to an immunogenic protein from *Brucella suis* (UniProt AC# P0A3U9), whereas Pfam indicates it could be related to the DNA repair protein REV1. Neither c6313_g1_i1.p1 nor c1505_g2_i1.p1 matched any proteins from the Blast homology or Pfam protein families. Consequently, the nature, structure, and function of the three highly abundant proteins remains unknown.

**Table 3:** Summary of verified proteins following parameter optimization, manual inspection, and filtering (Stage 3) for *E. denticulatum* extracts A and B using both tryptic and semi-tryptic analysis. For each identified protein, the molecular weight, number of identified peptides, sequence coverage, protein score, riBAQ, I^L^_rel_, rTPM, subcellular localization, and localization probability. ^1^Subcellular localization and localization probability was computed using DeepLoc (Almagro Armenteros et al., 2017).

| Protein ID | MW<br>[kDa] | #Pep<br>A<br>tryp | #Pep<br>B<br>tryp | #Pep<br>A<br>semi | #Pep<br>B<br>semi | Seq.<br>cov. A<br>tryp [%] | Seq.<br>cov. B<br>tryp<br>[%] | Seq.<br>cov. A<br>semi<br>[%] | Seq.<br>cov. B<br>semi<br>[%] | Score<br>tryp | Score<br>semi | riBAQ<br>A tryp | I<sub>rel</sub><sup>L</sup> A<br>tryp | I<sub>rel</sub><sup>L</sup> A<br>semi | riBAQ<br>B tryp | I<sub>rel</sub><sup>L</sup> B<br>tryp | I<sub>rel</sub><sup>L</sup> B<br>semi | rTPM | Subcellular<br>localization<sup>1</sup> | Subcell<br>score<sup>1</sup> |
| --- | --- | --- | --- | --- | --- | --- | --- | --- | --- | --- | --- | --- | --- | --- | --- | --- | --- | --- | --- | --- |
| c6313_g1_i1.p1 | 21.153 | 7 | 1 | 13 | 5 | 36.3 | 7.4 | 47.9 | 17.4 | 323.3 | 323.3 | 32.3% | 35.5% | **32.1%** | 3.4% | 3.8% | **23.3%** | 0.29% | Extracellular | 0.6985 |
| c7052_g1_i1.p1 | 24.213 | 7 | 8 | 16 | 7 | 36.1 | 34.8 | 45.4 | 31.7 | 323.3 | 323.3 | 31.9% | 35.2% | **33.0%** | 31.9% | 35.9% | **25.1%** | 0.02% | Extracellular | 0.9128 |
| c1505_g2_i1.p1 | 15.778 | 4 | 4 | 7 | 5 | 30 | 30 | 30 | 30 | 323.3 | 323.3 | 15.1% | 11.3% | **17.1%** | 33.4% | 25.4% | **28.0%** | 0.14% | Extracellular | 0.4441 |
| c4354_g1_i1.p1 | 40.332 | 8 | 1 | 7 | 3 | 24.6 | 3.5 | 21.7 | 5.6 | 323.3 | 323.3 | 5.2% | 4.3% | **4.0%** | 0.2% | 0.1% | **0.3%** | 0.15% | Extracellular | 0.8483 |
| c7052_g1_i2.p1 | 23.965 | 6 | 5 | 15 | 5 | 30.8 | 25.1 | 40.1 | 25.1 | 163.3 | 140.2 | 4.9% | 4.5% | **4.1%** | 8.3% | 7.8% | **6.8%** | 0.06% | Extracellular | 0.9431 |
| c17304_g1_i1.p1 | 27.965 | 6 | 2 | 6 | 1 | 23 | 6.7 | 19.7 | 3 | 188.3 | 190.4 | 2.9% | 3.4% | **3.2%** | 4.8% | 5.7% | **3.5%** | 1.08% | Extracellular | 0.5089 |
| c8421_g1_i1.p1 | 59.681 | 4 | 0 | 4 | 1 | 10 | 0 | 10 | 2.3 | 323.3 | 323.3 | 2.6% | 1.6% | **1.6%** | 0.0% | 0.0% | **0.0%** | 0.04% | Membrane | 0.9998 |
| c6458_g1_i1.p1 | 46.381 | 3 | 0 | 7 | 0 | 10.8 | 0 | 17.5 | 0 | 303.3 | 145.4 | 2.2% | 1.4% | **1.8%** | 0.0% | 0.0% | **0.0%** | 0.12% | Extracellular | 0.8601 |
| c5232_g1_i1.p1 | 18.952 | 2 | 0 | 2 | 0 | 13.5 | 0 | 13.5 | 0 | 125.3 | 104.9 | 1.2% | 1.1% | **1.1%** | 0.0% | 0.0% | **0.0%** | 0.03% | Extracellular | 0.6419 |
| c4671_g1_i1.p2 | 29.874 | 4 | 0 | 3 | 0 | 19.9 | 0 | 17 | 0 | 52.5 | 31.4 | 0.5% | 0.4% | **0.2%** | 0.0% | 0.0% | **0.0%** | 0.03% | Plastid | 0.995 |
| c7216_g1_i1.p1 | 25.446 | 2 | 0 | 3 | 0 | 10.8 | 0 | 16.9 | 0 | 16.1 | 19.3 | 0.4% | 0.3% | **0.4%** | 0.0% | 0.0% | **0.0%** | 0.08% | Extracellular | 0.8121 |
| c17615_g1_i1.p1 | 27.973 | 2 | 0 | 0 | 0 | 7.7 | 0 | 0 | 0 | 13.9 | 0.0 | 0.2% | 0.2% | **0.0%** | 0.0% | 0.0% | **0.0%** | 0.06% | Extracellular | 0.9751 |
| c6963_g2_i1.p1 | 165.47 | 10 | 2 | 4 | 2 | 6.5 | 1.6 | 2.1 | 1.6 | 108.7 | 55.5 | 0.1% | 0.1% | **0.1%** | 0.5% | 0.5% | **0.6%** | 0.04% | Plastid | 0.6933 |
| c4090_g1_i1.p1 | 16.129 | 1 | 2 | 1 | 1 | 6.8 | 12.9 | 6.8 | 6.1 | 15.3 | 11.2 | 0.1% | 0.1% | **0.1%** | 0.5% | 0.7% | **0.0%** | 0.02% | Plastid | 0.9815 |
| c231_g1_i1.p1 | 18.006 | 2 | 0 | 0 | 0 | 10.2 | 0 | 0 | 0 | 11.7 | 0.0 | 0.1% | 0.1% | **0.0%** | 0.0% | 0.0% | **0.0%** | 0.05% | Extracellular | 0.9998 |
| c2364_g1_i1.p1 | 50.492 | 1 | 5 | 2 | 5 | 2.4 | 9.9 | 4.1 | 9.9 | 135.4 | 105.1 | 0.1% | 0.1% | **0.1%** | 8.4% | 10.8% | **6.9%** | 0.23% | Cytoplasm | 0.7655 |
| c926_g1_i1.p1 | 79.764 | 3 | 0 | 2 | 0 | 3.6 | 0 | 2.6 | 0 | 20.2 | 10.9 | 0.1% | 0.1% | **0.1%** | 0.0% | 0.0% | **0.0%** | 0.04% | Extracellular | 0.9924 |
| c6373_g1_i1.p1 | 119.64 | 4 | 1 | 2 | 0 | 4.1 | 0.9 | 2.3 | 0 | 79.2 | 40.8 | 0.1% | 0.0% | **0.0%** | 0.1% | 0.1% | **0.0%** | 0.06% | Extracellular | 0.5704 |
| c6656_g1_i1.p1 | 43.007 | 3 | 3 | 1 | 3 | 8.8 | 8.5 | 3.3 | 8.5 | 79.4 | 75.8 | 0.0% | 0.0% | **0.0%** | 7.0% | 7.4% | **5.4%** | 0.12% | Plastid | 0.9982 |
| c6834_g1_i1.p3 | 22.388 | 1 | 1 | 0 | 0 | 5.3 | 6.7 | 0 | 0 | 15.3 | 0.0 | 0.0% | 0.0% | **0.0%** | 0.5% | 0.7% | **0.0%** | 0.06% | Plastid | 0.9985 |
| c2556_g1_i1.p1 | 57.477 | 1 | 3 | 0 | 0 | 2 | 4.9 | 0 | 0 | 19.4 | 0.0 | 0.0% | 0.0% | **0.0%** | 0.8% | 0.8% | **0.0%** | 0.05% | Plastid | 0.567 |
| c1613_g1_i1.p1 | 32.099 | 0 | 2 | 1 | 2 | 0 | 7.8 | 5.4 | 9.5 | 35.0 | 31.5 | 0.0% | 0.0% | **0.0%** | 0.4% | 0.4% | **0.2%** | 0.09% | Plastid | 0.9671 |
| c3760_g1_i1.p1 | 32.533 | 0 | 0 | 3 | 0 | 0 | 0 | 6.6 | 0 | 0.0 | 253.1 | 0.0% | 0.0% | **0.0%** | 0.0% | 1.0% | **0.0%** | 0.06% | Lysosome | 0.3775 |

**Figure 2:**
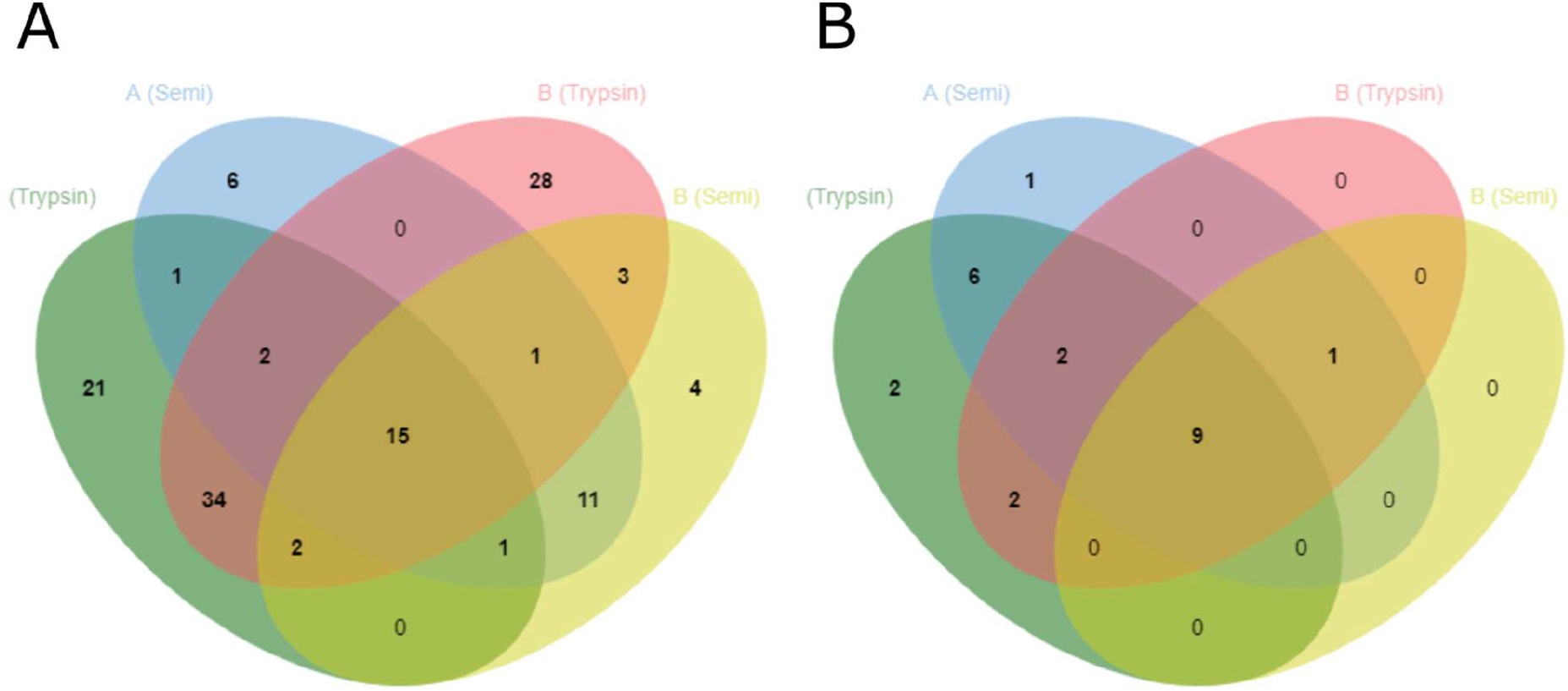
4-way Venn diagrams showing identified peptides (A) and proteins (B) with optimized parameters (5% FDR and minimum score threshold) and following filtering for extract A using tryptic analysis (green), extract A using semi-tryptic analysis (blue), extract B using tryptic analysis (red), and extract B using semi-tryptic analysis (yellow). List sizes (in the same order) for peptides (A) are 76, 37, 85, and 37 for a total of 129 identified peptides. List sizes (in the same order) for proteins (B) are 21, 19, 14, and 10 for a total of 23 identified proteins.

**Figure 3:**
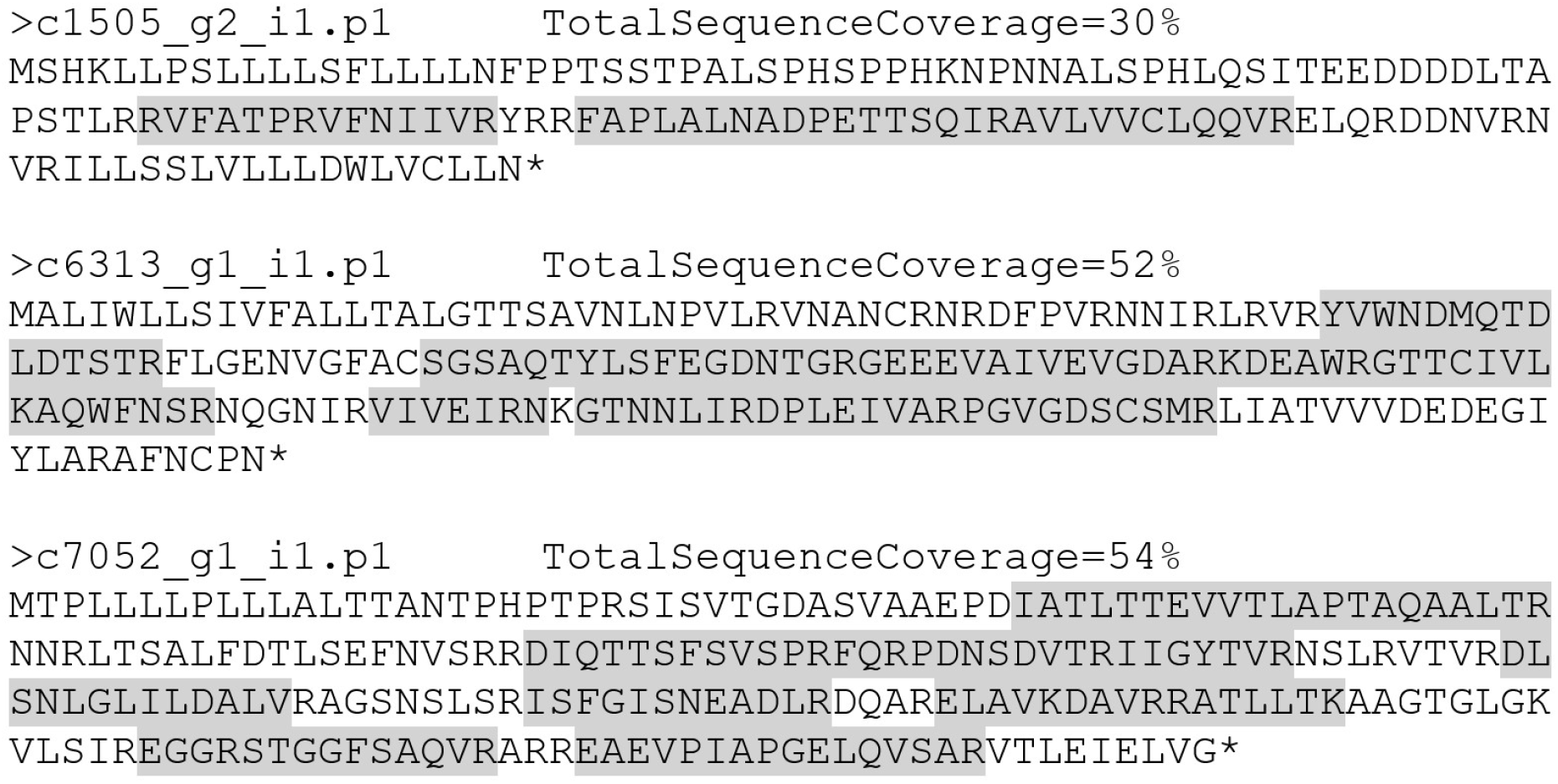
Protein sequence and experimental sequence coverage across both extracts and analysis methods (highlighted in grey) for the three most abundant *E. denticulatum* proteins identified. All three proteins passed final selection criteria (Stage 3) and accounted for 82.2% and 76.4% (quantified by I^L^_rel_ using semi-specific analysis) of the verified, seaweed-specific proteins in extracts A and B, respectively. Including the isoform of c7052_g1_i1 (c7052_g1_i1 – not shown), the proteins account for and 86.4% and 83.2%, respectively.

Filtering resulted in improved correlation between the two extracts up to a PCC of 0.91 for relative abundances quantified by I^L^_rel_ (Figure 4). This indicates that in light of all the complications, the two protein extracts are in fact comparable, when all redundancy and contamination was addressed. Furthermore, the in-sample correlation between riBAQ and I^L^_rel_ (PCC 0.87-1.0) indicated that I^L^_rel_ may in fact be quite powerful analogue to riBAQ for non-standard (i.e. semi- or unspecific) analysis. As semi-specific in most cases increase both number of identified peptides as well as the sequence coverage on the individual protein level I^L^_rel_ could be a powerful tool in the analysis of proteins where partial (non-specific) hydrolysis is observed, as this will include all peptide originating from the parent proteins rather than proteotryptic peptides alone.

**Figure 4:**
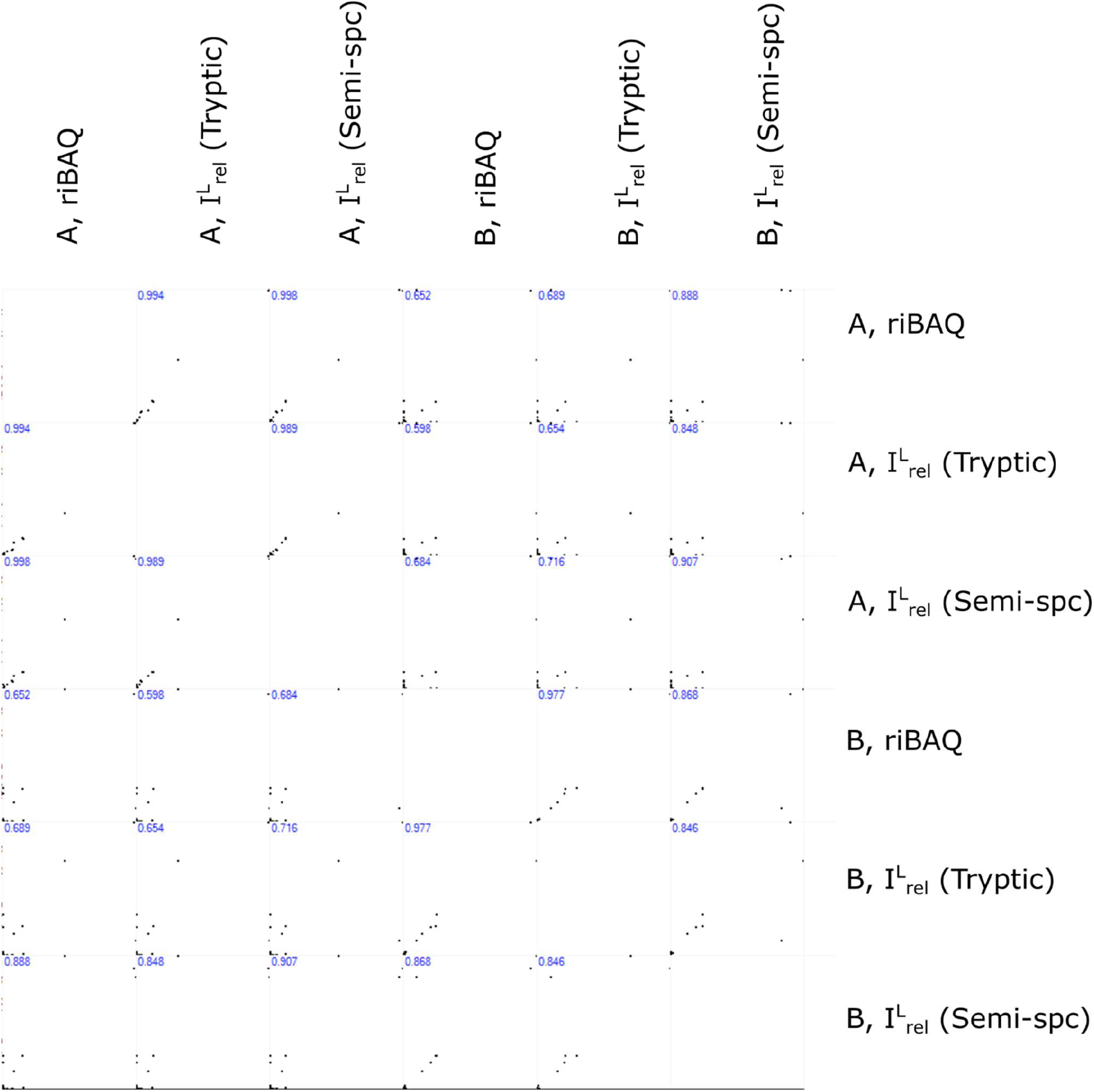
Correlation of relative protein abundances between extracts (A and B), analysis conditions (tryptic and semi-specifc), and quantification method (riBAQ and I^L^_rel_) following manual validation, filtering, and re-quantification (Stage 3). Pearson Correlation Coefficients are shown in blue in the upper left corner of each sub-plot.

Considering the level of contamination in the extracts as outlined above, this naturally affects the potential yield in targeted processing of the proteins. Including all initially identified peptides/protein including the common contaminants, the final list of quantified proteins constitute 78% of the total protein for extract A but merely 6.0% of the total protein for extract B. This correspond to the verified *E. denticulatum* proteins (Stage 3) constituting 5.6% and 4.2% of the total extract mass, based on the total protein content for the individual extracts. The observed level of contamination also indicated that although the total protein content was significantly increased in extract B, this may also come at a high cost in terms of applicability. However, as protein contamination can occur at all stages from processing facility to analysis lab, this should be investigated further. Furthermore, the tryptic analysis showed a significantly lower number of peptides and relative abundance for c6313_g1_i1.p1 compared to the semi-tryptic analysis for extract B. This could indicate that this particular protein is subject to partial hydrolysis during the additional processing, which again strengthens the use of the semi-specific analysis for this type of protein extract. The high degree of exogenous protein identified in extract B may also explain why the N-to-protein conversion factor of 5 appears to give much better results for extract A, and why extract B appears to be more accurately estimated using the Jones factor of 6.25 (Table 1).

### 3.4. Enrichment of extracellular proteins

In Figure 5, the relative subcellular distribution of proteins predicted by DeepLoc, is presented. For the transcriptome analysis (Fig 5C), a relative broad distribution of proteins (by rTPM) is observed with the majority of proteins being ascribed to the nucleus (24%), plastid (22%), cytoplasm (20%), mitochondria (18%), and extracellular (7%). This distribution does not correlate with the protein distribution established by LC-MS/MS, regardless of data analysis conditions employed. In fact, there is a very significant enrichment in extracellular proteins. For extract A (Fig 5A), almost exclusively extracellular proteins are identified (97%) by I^L^_rel_,. While extract B (Fig 5B) has some content of plastid and cytoplasmic protein, the majority of identified proteins are extracellular (87%) by I^L^. The three primary proteins in both extracts are all classified as being extracellular. Although c7052_g1_i1.p1 shows homology to a periplasmic proteins by BlastP, it has a very high extracellular localization probability using DeepLoc (Table 2). At the individual protein level, the extracellular protein with the highest rTPM of 1.1%, c17304_g1_i1.p1 (see Table A.3), was determined to constitute 3.2-3.5% of the molar protein content. Although still significantly abundant, the three highly abundant extracellular proteins described above (I^L^_rel_ 17-33% each) merely constituted 0.02-0.29% on the transcript level, indicating that the extraction method is not selective for extracellular proteins per se, but rather a few selected extracellular proteins.

**Figure 5:**
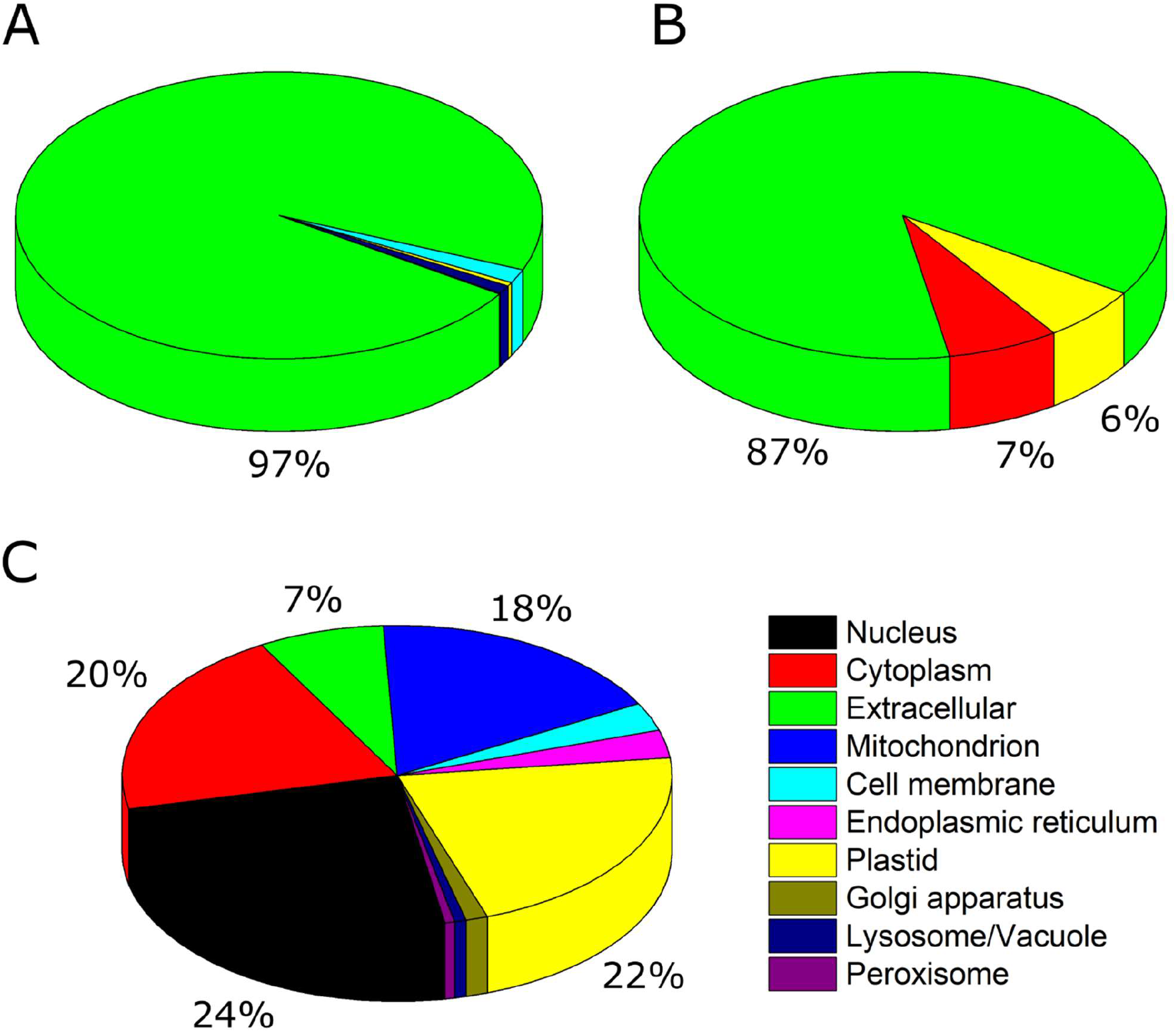
Relative subcellular protein distribution as predicted by DeepLoc (Almagro Armenteros et al., 2017) for A: Protein extract A. B: Protein extract B. For both protein extracts, relative abundance was estimated by I^L^_rel_ through semi-tryptic analysis using optimized parameters, following manual inspection, validation, and filtering. C: Transcriptome analysis (by rTPM).

The fact that extracellular protein were almost exclusively identified in the extracts, is also very likely to explain the low extraction yields observed at the pilot plant (unpublished data from CP Kelco). From 20 kg of seaweed, 155 g material was obtained using a 1000 L extraction tank (Extract A). The protein content (by Kjeldahl-N and converted using the Jones factor) of 7.1% correspond to merely 11 g of protein following extraction corresponding to a protein yield of 0.055%. Further processing to concentrate protein by acid precipitation (Extract B) yielded 6.7 g of product with 71% protein corresponding to 4.8 g of protein and consequently a loss of 57% protein mass and thus an even lower yield (0.024%). These findings indicate that the hot-water extraction used to obtain the extracts, is not capable, to a significant degree, to disrupt cells and release intracellular proteins. Low protein yields using simple aqueous extraction from *E. denticulatum* has previously been reported in literature (Bjarnadóttir et al., 2018; Fleurence, Le Coeur, Mabeau, Maurice, & Landrein, 1995). This, in turn, implies that there is still a significant potential for protein extraction from the seaweed and other approaches such as for instance pressurized and supercritical fluid extraction (Herrero, Sánchez-Camargo, Cifuentes, & Ibáñez, 2015), addition of cofactors (Harnedy & FitzGerald, 2013a; Maehre, Edvinsen, Eilertsen, & Elvevoll, 2016), microwave-assisted extraction (Magnusson et al., 2019), ultrasound-assisted extraction (Bleakley & Hayes, 2017), or any combination thereof (Cermeño, Kleekayai, Amigo-Benavent, Harnedy-Rothwell, & FitzGerald, 2020), may be more suitable. Enzyme assisted extraction (EAE) is an emerging technology for seaweed protein extraction, showing great potential (Hardouin et al., 2016; Naseri, Marinho, Holdt, Bartela, & Jacobsen, 2020; Terme et al., 2020; Vásquez, Martínez, & Bernal, 2019). In a recent study, enzyme assisted extraction of *E. denticulatum* increased the protein yield up to 60% using Alcalase^®^ or Viscozyme^®^ (0.2% w/w) at pH 7 and room temperature (Naseri, Jacobsen, et al., 2020). The increased protein extraction efficiency was furthermore obtained without compromising the downstream carrageenan production. However, this method is not at present implemented by the carrageenan industry.

## 4. Conclusion

Using *de* novo transcriptome assembly, we were able to construct a novel reference proteome for *E. denticulatum*, which was used to characterize two pilot-scale, hot-water extracts. Although further processing (extract B) increased protein content significantly (compared to extract A), the aqueous solubility of both was quite low and both extracts displayed a high degree of smear and a lack of distinct protein bands by SDS-PAGE. A slightly alkaline pH and addition of a small amount of detergent fully solubilized the protein. From proteomics studies, using label-free quantification of non-standard protein digests via a novel length-normalized relative abundance approach, we determined that further extract processing may have introduced a significant amount of contaminant proteins not originating from the seaweed. After filtering of contaminant proteins and potential false-positive protein identifications, the protein content from the two extracts correlated quite well. Using subcellular localization prediction, we determined that both extracts were highly enriched in extracellular protein compared to the expected protein distribution from quantitative transcriptome analysis and estimated protein copy number. In fact, more than 75% of the seaweed-specific protein identified and quantified, was constituted by merely three proteins, which were predicted to be extracellular. Extracellular protein enrichment indicates that hot-water extraction is not capable of extracting intracellular proteins, but may be useful for isolation of extracellular protein content on large, industrial scale. Further processing of seaweed extracts is useful for increasing total protein content, but it requires further optimization to reduce the introduction of a large degree of exogenous protein, the depletion of species-specific proteins, and the significant loss in total protein. Nevertheless, this study illustrates the applicability of quantitative proteomics for characterization of extracts to be used as potential sources of novel food protein or bioactive peptides. Furthermore, the results clearly demonstrate the power of the methodology, particularly in combination with quantitative transcriptomics and bioinformatics, for evaluating extraction methods and for use as a guide in the development and optimization of industrial processes.

## 5. Acknowledgements

The authors would like to thank CP Kelco and in particular, Senior Scientist Jimmy Sejberg for supplying the seaweed protein extracts and for fruitful discussions related to analysis and manuscript preparation.

## 6. Funding

This work was supported by Innovation Fund Denmark (Grant number 7045-00021B (PROVIDE)).

## 7. Author Contribution

S.G.: Conceptualization, Methodology, Formal analysis, Investigation, Writing – original draft preparation, Writing – review and editing, Visualization. M.P.: Methodology, Formal analysis, Investigation, Writing – original draft preparation, Writing – review and editing. P.M.: Conceptualization, Methodology, Writing – review and editing, Supervision. S.L.H.: Writing – original draft preparation, Writing – review and editing. C.J.: Writing – original draft preparation, Writing – review and editing, Funding acquisition. P.J.G-M.: Methodology, Writing – review and editing. E.B.H.: Conceptualization, Writing – review and editing, Project administration, Funding acquisition. M.T.O.: Conceptualization, Writing – review and editing, Supervision.

## 8. Conflict of Interest

The authors declare no conflict of interest.

## 9. Data Availability

MaxQuant output files (txt folder) can be accessed through the linked Mendeley Data repository (Gregersen, 2020). Raw MS data can be made available upon request.

## Notes

http://dx.doi.org/10.17632/y4kmnb3tvx.1

### Competing Interest Statement

The authors have declared no competing interest.

