## Supplementary Information for "Proteomic characterization of pilot scale hot-water extracts from the industrial carrageenan red seaweed *Eucheuma denticulatum*"

*Peptide identification and validation*

Figure A.8 (left) shows a 4-way Venn diagram of peptide level identifications, using both standard tryptic digestion and unspecific digestion for the two enzymatic extracts with 1% false discovery rate (FDR) (i.e. initial search). From here, we see that we identified 279 unique peptides across the two extracts and conditions (Table A.8). Surprisingly, very few were shared between all four (21) while more than half (153) of the peptides were unique to one extract and analysis condition. Additionally, almost half (119) were exclusive to either one or both tryptic analyses while only half of that (60) were exclusive to either one or both unspecific analyses. This trend is also in line with the observation that analysis using *in silico* tryptic digestions generally appears to result in a higher number of peptide identifications. As unspecific searches uses both a significantly larger search space, is also employs a very large decoy database for FDR-control. When tryptic restrictions to peptide termini are also lifted from the decoy database, this dramatically increases the chance of identifying a non-contaminant decoy peptide by chance (Chi, Liu, Yang, Zeng, Wu, Zhou, Niu, et al., 2018; Chi, Liu, Yang, Zeng, Wu, Zhou, Wang, et al., 2018). This is also reflected in many of the identified tryptic peptides having low scores and significant posterior error probabilities (PEP). Indeed, by increasing peptide-level (and protein-level) FDR to first 5% (Figure A.9, B) and later 10% (Figure A.9, C), we were able increase the number of shared peptides while decreasing the number exclusively tryptic peptides (Table A.8).

A direct consequence of increasing FDR for unspecific searches to increase the number of shared peptides will be increasing the number of non-tryptic peptides. This was also observed here, as the number of exclusively unspecific peptides increased from 60 at 1% FDR over 74 at 5% FDR to 109 at 10% FDR. Although increasing peptide level FDR significantly from 1% to boost identification rates is generally not regarded the best approach (Jones, Siepen, Hubbard, & Paton, 2009; Mao et al., 2019), optimizing the peptide level PDR is pivotal for label-free quantification (Gupta & Pevzner, 2009; Lu, Vogel, Wang, Yao, & Marcotte, 2007). The overall cost in the unspecific analysis is not enormous, as peptide scores and PEP values are still better than those observed for tryptic analysis with 1% FDR, implying much better peptide spectrum matches (PSMs). To accommodate the generally low overlap between conditions, we also employed semi-specific tryptic restrictions to the target and decoy databases using both 1% FDR (Figure A.9, D) and 5% FDR (Figure A.9, E). As expected, this approach significantly increased the number of shared peptides with the tryptic *in silico* digest compared to the unspecific searches at the same FDR (Table A.8).

On the protein level, unspecific analysis at 1% FDR from the initial searches provided a significant amount of proteins identified in either extract A or both extracts by *in silico* tryptic digestions, whereas very few were identified exclusively for unspecific analysis on extract A. No proteins were identified exclusively for extract B in either approach (Figure A.8, right). This is a direct consequence of the increased search space for unspecific analysis described above. Similarly to the peptide level observations, increasing unspecific FDR to 5% (Figure A.10, B) or 10% (Figure A.10, C) increases the number of identified proteins by this approach, but also decreases the number of shared proteins. Furthermore, the number of proteins exclusive for extract B increase dramatically form none observed at 1% FDR (Figure A.10, A). Employing semi-specific *in silico* digestion somewhat alleviates this at both 1% FDR (Figure A.10, D) and 5% FDR (Figure A.10, E). In particular for the semi-specific analysis at 5% FDR, there is a good correlation on the number of proteins identified across both extracts and conditions.

*Filtration of optimized protein list.*

Using the optimized analysis parameters, we identified a total of 40 proteins across samples and conditions following contaminant filtering (Tables S5 and S6). Nevertheless, the identified and quantified proteins were subjected to manual validation and filtering in order to remove low confidence (i.e. potential false positive) identifications and potential contaminants introduced during sample processing, that were not identified using the “common contaminants” list in MaxQuant (Jurgen Cox et al., 2009). In short, seven quality-based filtering criteria were established, and proteins falling below the threshold in four of five criteria were subsequently filtered prior to final quantification. The full list of proteins and the adherence to the filtering criteria can be seen in Table A.7. The criteria were defined as:

- Low peptide ID count: An identified protein is only identified by two unique peptides across samples and conditions.
- All high PEP: All identified peptides are identified by low quality PSMs as evaluated by a PEP > 0.005. Proteins, where > 50% of the identified peptides are based on a PEP > 0.005 are listed in parenthesis.
- Low protein score: The highest protein score obtained is < 40.
- Low sequence coverage: Protein sequence coverage < 5%. Proteins with sequence coverage between 5% and 10% are listed in parenthesis.
- Low MS/MS count: The protein is only associated with ≤ 5 MS/MS scans across samples and conditions.
- Not determined in extract A: As Extract B is a further processing of extract A, proteins only identified in B is included here.
- Highly enriched in B: As Extract B is a further processing of extract A, protein that are > 10-fold enriched (relative) in B are included here.

*Evaluation of length-normalized relative intensity for label-free quantification of non-standard protein digests*

During the initial iterative process, the simple approach of merely applying relative intensity $I_{rel}$, resulted in good correlation with rIBAQ for extract A (PCC > 0.97), as seen in Figure A.4. However, the PCCs for extract B was much worse and as low as 0.049 between iBAQ and $I_{rel}$ for unspecific digestion at 1% FDR (Figure

A.5). Normalizing the protein intensities to their length before calculating the relative abundance, $I^L_{rel}$, increased correlation with riBAQ for both extracts (Figure A.6; A.7). For extract A, the PCC increased to 0.99 or higher while for extract B, the correlation between iBAQ and $I^L_{rel}$ for unspecific digestion at 1% FDR was still very low at 0.15. As expected, significantly higher correlation in quantification is observed when adjusting FDR within a specific *in silico* digestion approach (e.g. between all unspecific digestions with varying FDR). Nevertheless, using $I^L_{rel}$ for relative protein quantification appear highly feasible based on extract A. iBAQ normalizes by theoretical tryptic peptides from *in silico* digestion and is thus proportional to molar quantities of proteins. As no (sequence dependent) restrictions on theoretical peptides exist for unspecific searches (besides minimum and maximum length), the number of theoretical peptides is governed by the protein length. By normalizing to length on the individual protein level, $I^L_{rel}$ may be regarded as the unspecific analogue to riBAQ. In fact, $I^L_{rel}$ may, to some extent, also be regarded as an analougue to the total protein approach (TPA) originally described by Wiśniewski (Wiśniewski, 2017; Wiśniewski et al., 2012; Wiśniewski, Wegler, & Artursson, 2018). In TPA, the total protein MS1 intensity is normalized to the sum product of all protein intensities and their molecular weight, where we instead use protein length. The two quantitative metrics should overall correlate quite well, but this is currently under investigation

Unspecific digestion introduces a very large degree of randomness in quantification and increases the risk of false positive (or contaminant peptide) identifications by increasing size and complexity of the reference database (Swaney, Wenger, & Coon, 2010). If the peptide is furthermore modified, this has been shown to be associated with higher scores in Andromeda, which poses a significant problem in quantification (Bogdanow, Zauber, & Selbach, 2016). The amount of modified detected peptides in this study was quite low, and hence, this is more of a general consideration rather that applicable in this specific case. Some of the peptides identified using unspecific and semi-tryptic digestion may also be a direct result of non-optimized experimental conditions for tryptic digestion, leading to nonspecific cleavage by trypsin (Fang et al., 2015). But as the extracts are digested with trypsin, we ultimately considered the approach redundant and decided to only apply specific and semi-specific tryptic digestion for the *in silico* PSM DB. Although increasing correlation of riBAQ with $I^{L}_{rel}$ from unspecific digests compared to semi-specific digests, in particular for extract B (Figure A.7), this is largely governed by a few, low scoring protein identifications of substantial abundance, resulting in lower PCCs for semi-specific analysis.

The set of optimized parameters applied for quantification differ somewhat from the parameters applied in conventional proteomics studies, and are a direct consequence of the low extract quality. Indeed, using standard tryptic analysis, merely 219 peptides were identified (Table A.8), where proteomics studies often result in identification of thousands of peptides identified. For instance, in a similar study of protein extracts from the potato processing industry, we identified more over 1000 peptides in the tryptic digest of the final waste stream, Protamylasse (i.e. the lowest quality sample), and almost 8000 peptides for the untreated potato fruit juice under the same analysis conditions (García-Moreno et al., 2020). Due to the low number of peptide (and protein) IDs, not only is the FDR of utmost importance, but the risk of quantifying proteins based on low peptide and protein scores is obvious. In addition, peptide score in MaxQuant is not only based on PEP, but also by the intensity of a given feature (Jürgen Cox et al., 2011; Tiwary et al., 2019; Tyanova, Temu, & Cox, 2016). Consequently, high intensity features with significant PEP (i.e. potential false positives), which in other studies may have been filtered out, will obtain a sufficiently high peptide score and be used in protein quantification. Ultimately, this leads to false identification of proteins with a significant relative abundance, which impairs further analysis. By applying more stringent thresholds on both peptide and protein level, this is alleviated to some extent. Nevertheless, it may be needed to inspect and evaluate PEPs rather than apply threshold filtering on peptide score alone, as PEP relies solely on PSM and sequence-dependent features (Jürgen Cox et al., 2011; Tiwary et al., 2019; Tyanova et al., 2016). Ultimately, this relies on sample quality and analytical approach.

In order to obtain full justification for the quantitative approach, the metric must be applied to a sample that has undergone truly unspecific digestion. Further validation of the metric is needed by direct comparison of fully digested tryptic samples and samples digested with unspecific protease – potentially for a limited time to prevent overdigestion and full hydrolysis. For comparison, it would be highly beneficial to include proteases with other specificities as well. Comparison of DDA and DIA is also needed, as the stochastic sampling of precursor ions in DDA may be somewhat biased (Van Puyvelde et al., 2020). Ultimately, the approach should be benchmarked against conventional riBAQ and TPA.

| | |
|---|---|
| Total trinity 'genes' | 19248 |
| Total trinity transcripts | 21444 |
| % GC | 52.73 |
| Stats based on ALL transcript contigs | |
| Contig N10 | 4644 |
| Contig N20 | 3586 |
| Contig N30 | 2920 |
| Contig N40 | 2385 |
| Contig N50 | 1958 |
| Median contig length | 507 |
| Average contig | 1021.23 |
| Total assembled bases | 21899322 |
| Stats based on ONLY LONGEST ISOFORM per 'GENE' | |
| Contig N10 | 4589 |
| Contig N20 | 3506 |
| Contig N30 | 2852 |
| Contig N40 | 2316 |
| Contig N50 | 1891 |
| Median contig length | 477 |
| Average contig | 977.18 |
| Total assembled bases | 18808710 |

Table A.1: Basic contig statistic of the assembled *de novo* transcriptome of *E. denticulatum* using Trimmomatic and Trinity.

| |
|---|
| 13394021 reads; of these: |
| 13394021(100%) were paired; of these: |
| 533933 (3.99%) aligned concordantly 0 times |
| 11232960 (83.87%) aligned concordantly exactly 1 time |
| 1627128 (12.15%) aligned concordatnly >1 times |
| 533933 (3.99%) aligned concordantly 0 times; of these |
| 170199 (31.88%) aligned discordantly 1 time |
| 363734 pairs aligned 0 times concordantly or discordantly; of these: |
| 727468 mates make up pairs; of these: |
| 480776 (66.09%) aligned 0 times |
| 110369 (15.17%) aligned exactly 1 time |
| 136323 (18.74%) aligned >1 times |
| **98.21% overall alignment rate** |

Table A.2: Percentage of the remapped reads to the assembly by Bowtie2 for the *de novo* transcriptome assembly of *E. denticulatum* using Trinity.

Table A.3: A full list of protein accessions and their associated TPMs, rTPMs, Pfam functions, BlastX targets, BlastP targets, predicted subcellular localization, and localization probability (table appended separately as supplementary .xlsx file)

Table A.4: Reference proteome as constructed by *de novo* transcriptome assembly (appended separately as supplementary .fasta file)

| Protein ID | Mol. weight [kDa] | Sequence length | Number of proteins tryp | Peptides A tryp | Peptides B tryp | Unique peptides A tryp | Unique peptides B tryp | Sequence coverage A tryp [%] | Sequence coverage B tryp [%] | Score tryp | MS/MS count A tryp | MS/MS count B tryp | riBAQ A tryp | riBAQ B tryp | rI_L A tryp | rI_L B tryp |
|---|---|---|---|---|---|---|---|---|---|---|---|---|---|---|---|---|
| c10492_g1_i1.p1 | 40.34 | 369 | 1 | 0 | 2 | 0 | 2 | 0 | 4.3 | 12.389 | 0 | 10 | 0.0% | 1.2% | 0.0% | 1.1% |
| c1294_g1_i1.p1 | 37.606 | 341 | 1 | 1 | 1 | 1 | 1 | 2.6 | 2.1 | 11.519 | 1 | 0 | 0.0% | 0.1% | 0.0% | 0.1% |
| c1357_g1_i1.p1 | 28.81 | 275 | 1 | 0 | 2 | 0 | 2 | 0 | 3.3 | 11.595 | 0 | 2 | 0.0% | 0.6% | 0.0% | 0.5% |
| c1505_g2_i1.p1 | 15.778 | 140 | 1 | 4 | 4 | 4 | 4 | 30 | 30 | 323.31 | 39 | 9 | 14.9% | 21.5% | 11.0% | 13.7% |
| c1613_g1_i1.p1 | 32.099 | 296 | 1 | 0 | 2 | 0 | 2 | 0 | 7.8 | 35.038 | 0 | 6 | 0.0% | 0.3% | 0.0% | 0.2% |
| c17161_g1_i1.p1 | 33.439 | 292 | 1 | 0 | 2 | 0 | 2 | 0 | 5.5 | 14.588 | 0 | 3 | 0.0% | 0.6% | 0.0% | 0.6% |
| c17201_g1_i1.p1 | 75.477 | 707 | 1 | 1 | 1 | 1 | 1 | 1.3 | 1.1 | 13.91 | 1 | 1 | 0.0% | 0.0% | 0.0% | 0.0% |
| c17231_g1_i1.p1 | 71.408 | 656 | 1 | 1 | 2 | 1 | 2 | 1.7 | 2.4 | 18.867 | 1 | 7 | 0.0% | 3.8% | 0.0% | 3.8% |
| c17304_g1_i1.p1 | 27.965 | 269 | 1 | 6 | 2 | 5 | 2 | 23 | 6.7 | 188.33 | 45 | 5 | 2.9% | 3.1% | 3.3% | 3.0% |
| c17615_g1_i1.p1 | 27.973 | 246 | 1 | 2 | 0 | 2 | 0 | 7.7 | 0 | 13.895 | 7 | 0 | 0.2% | 0.0% | 0.2% | 0.0% |
| c231_g1_i1.p1 | 18.006 | 166 | 1 | 2 | 0 | 2 | 0 | 10.2 | 0 | 11.702 | 3 | 0 | 0.1% | 0.0% | 0.1% | 0.0% |
| c2364_g1_i1.p1 | 50.492 | 464 | 1 | 1 | 5 | 1 | 5 | 2.4 | 9.9 | 135.35 | 6 | 19 | 0.1% | 5.4% | 0.1% | 5.8% |
| c2556_g1_i1.p1 c46_g1_i1.p1 | 57.477 | 510 | 2 | 1 | 3 | 1 | 3 | 2 | 4.9 | 19.385 | 1 | 4 | 0.0% | 0.5% | 0.0% | 0.4% |
| c3249_g1_i1.p1 | 24.109 | 219 | 1 | 2 | 0 | 2 | 0 | 8.7 | 0 | 11.121 | 2 | 0 | 0.0% | 0.0% | 0.1% | 0.0% |
| c4090_g1_i1.p1 | 16.129 | 147 | 1 | 1 | 2 | 1 | 2 | 6.8 | 12.9 | 15.342 | 2 | 2 | 0.1% | 0.3% | 0.1% | 0.4% |
| c4354_g1_i1.p1 | 40.332 | 374 | 1 | 8 | 1 | 8 | 1 | 24.6 | 3.5 | 323.31 | 56 | 1 | 5.1% | 0.1% | 4.2% | 0.1% |
| c4671_g1_i1.p2 | 29.874 | 271 | 1 | 4 | 0 | 4 | 0 | 19.9 | 0 | 52.513 | 8 | 0 | 0.5% | 0.0% | 0.4% | 0.0% |
| c4757_g1_i1.p1 | 38.191 | 338 | 1 | 0 | 2 | 0 | 2 | 0 | 5.3 | 18.758 | 0 | 5 | 0.0% | 1.7% | 0.0% | 1.7% |
| c4921_g1_i1.p1 | 38.874 | 344 | 1 | 0 | 3 | 0 | 3 | 0 | 8.7 | 19.707 | 0 | 3 | 0.0% | 0.6% | 0.0% | 0.6% |
| c5168_g1_i1.p1 | 25.142 | 230 | 1 | 0 | 2 | 0 | 2 | 0 | 10.4 | 21.494 | 0 | 3 | 0.0% | 0.2% | 0.0% | 0.3% |
| c5232_g1_i1.p1 | 18.952 | 178 | 1 | 2 | 0 | 2 | 0 | 13.5 | 0 | 125.25 | 10 | 0 | 1.2% | 0.0% | 1.1% | 0.0% |
| c6313_g1_i1.p1 | 21.153 | 190 | 1 | 7 | 1 | 7 | 1 | 36.3 | 7.4 | 323.31 | 169 | 9 | 31.9% | 2.2% | 34.8% | 2.1% |
| c6373_g1_i1.p1 | 119.64 | 1078 | 1 | 4 | 1 | 4 | 1 | 4.1 | 0.9 | 79.189 | 7 | 1 | 0.1% | 0.1% | 0.0% | 0.0% |
| c6458_g1_i1.p1 | 46.381 | 435 | 1 | 3 | 0 | 3 | 0 | 10.8 | 0 | 303.32 | 37 | 0 | 2.2% | 0.0% | 1.4% | 0.0% |
| c6656_g1_i1.p1 | 43.007 | 399 | 1 | 3 | 3 | 3 | 3 | 8.8 | 8.5 | 79.436 | 3 | 12 | 0.0% | 4.5% | 0.0% | 4.0% |
| c6797_g1_i1.p1 | 115.98 | 1048 | 1 | 0 | 2 | 0 | 2 | 0 | 1.7 | 15.61 | 0 | 3 | 0.0% | 0.1% | 0.0% | 0.1% |
| c6825_g1_i1.p3 c13559_g1_i1.p1 | 13.634 | 125 | 2 | 2 | 2 | 2 | 2 | 12.8 | 12.8 | 67.854 | 5 | 11 | 1.1% | 23.8% | 1.8% | 33.8% |
| c6834_g1_i1.p3 | 22.388 | 208 | 1 | 1 | 1 | 1 | 1 | 5.3 | 6.7 | 15.329 | 1 | 1 | 0.0% | 0.3% | 0.0% | 0.4% |
| c6945_g1_i1.p2 | 46.234 | 419 | 1 | 1 | 3 | 1 | 3 | 3.6 | 10.3 | 171.3 | 4 | 24 | 0.0% | 2.2% | 0.0% | 2.7% |
| c6963_g2_i1.p1 | 165.47 | 1543 | 1 | 10 | 2 | 10 | 2 | 6.5 | 1.6 | 108.74 | 23 | 11 | 0.1% | 0.3% | 0.1% | 0.3% |
| c7052_g1_i1.p1 | 24.213 | 227 | 1 | 7 | 8 | 2 | 5 | 36.1 | 34.8 | 323.31 | 145 | 36 | 31.6% | 20.6% | 34.6% | 19.4% |
| c7052_g1_i2.p1 | 23.965 | 227 | 1 | 6 | 5 | 1 | 2 | 30.8 | 25.1 | 163.32 | 18 | 13 | 4.9% | 5.3% | 4.4% | 4.2% |
| c7216_g1_i1.p1 | 25.446 | 231 | 1 | 2 | 0 | 2 | 0 | 10.8 | 0 | 16.086 | 5 | 0 | 0.4% | 0.0% | 0.3% | 0.0% |
| c8389_g1_i1.p1 | 21.023 | 194 | 1 | 1 | 1 | 1 | 1 | 4.1 | 4.1 | 14.912 | 1 | 1 | 0.0% | 0.7% | 0.0% | 0.8% |
| c8421_g1_i1.p1 | 59.681 | 570 | 1 | 4 | 0 | 4 | 0 | 10 | 0 | 323.31 | 63 | 0 | 2.6% | 0.0% | 1.6% | 0.0% |
| c926_g1_i1.p1 | 79.764 | 772 | 1 | 3 | 0 | 3 | 0 | 3.6 | 0 | 20.209 | 6 | 0 | 0.1% | 0.0% | 0.1% | 0.0% |

Table A.5: Summary of identifications and quantifications by tryptic analysis (5% FDR) following parameter optimization following filtering of common contaminants (Stage 2).

| Protein ID | Mol. weight [kDa] | Sequence length | Number of proteins semi | Peptides A semi | Peptides B semi | Unique peptides A semi | Unique peptides B semi | Sequence coverage A semi [%] | Sequence coverage B semi [%] | Score semi | MS/MS count A semi | MS/MS count B semi | rI_L A semi | rI_L B semi |
|---|---|---|---|---|---|---|---|---|---|---|---|---|---|---|
| c1275_g1_i1.p1 | 23.45 | 209 | 1 | 0 | 2 | 0 | 2 | 0 | 5.7 | 11.827 | 0 | 11 | 0.0% | 54.5% |
| c1505_g2_i1.p1 | 15.778 | 140 | 1 | 7 | 5 | 7 | 5 | 30 | 30 | 323.31 | 41 | 10 | 16.8% | 8.2% |
| c1613_g1_i1.p1 | 32.099 | 296 | 1 | 1 | 2 | 1 | 2 | 5.4 | 9.5 | 31.459 | 1 | 6 | 0.0% | 0.1% |
| c17231_g1_i1.p1 | 71.408 | 656 | 1 | 1 | 1 | 1 | 1 | 2.1 | 1.2 | 44.855 | 3 | 0 | 0.0% | 1.1% |
| c17304_g1_i1.p1 | 27.965 | 269 | 1 | 6 | 1 | 5 | 0 | 19.7 | 3 | 190.44 | 36 | 0 | 3.1% | 1.0% |
| c2364_g1_i1.p1 | 50.492 | 464 | 1 | 2 | 5 | 2 | 5 | 4.1 | 9.9 | 105.09 | 4 | 12 | 0.1% | 2.0% |
| c2788_g1_i1.p1 | 24.448 | 226 | 1 | 0 | 2 | 0 | 2 | 0 | 8.4 | 51.001 | 0 | 6 | 0.0% | 1.1% |
| c3760_g1_i1.p1 | 32.533 | 302 | 1 | 3 | 0 | 3 | 0 | 6.6 | 0 | 253.05 | 19 | 0 | 0.9% | 0.0% |
| c4090_g1_i1.p1 | 16.129 | 147 | 1 | 1 | 1 | 1 | 1 | 6.8 | 6.1 | 11.156 | 1 | 1 | 0.1% | 0.0% |
| c4354_g1_i1.p1 | 40.332 | 374 | 1 | 7 | 3 | 7 | 3 | 21.7 | 5.6 | 323.31 | 43 | 1 | 3.9% | 0.1% |
| c4671_g1_i1.p2 | 29.874 | 271 | 1 | 3 | 0 | 3 | 0 | 17 | 0 | 31.398 | 6 | 0 | 0.2% | 0.0% |
| c4757_g1_i1.p1 | 38.191 | 338 | 1 | 0 | 2 | 0 | 2 | 0 | 5.3 | 13.543 | 0 | 4 | 0.0% | 1.1% |
| c4921_g1_i1.p1 | 38.874 | 344 | 1 | 0 | 2 | 0 | 2 | 0 | 6.4 | 11.532 | 0 | 1 | 0.0% | 0.5% |
| c5168_g1_i1.p1 | 25.142 | 230 | 1 | 0 | 2 | 0 | 2 | 0 | 10.4 | 12.771 | 0 | 3 | 0.0% | 0.3% |
| c5232_g1_i1.p1 | 18.952 | 178 | 1 | 2 | 0 | 2 | 0 | 13.5 | 0 | 104.88 | 6 | 0 | 1.0% | 0.0% |
| c5952_g1_i1.p1<br>c1035_g1_i1.p1 | 41.634 | 374 | 2 | 0 | 2 | 0 | 2 | 0 | 8 | 12.249 | 0 | 7 | 0.0% | 0.4% |
| c6313_g1_i1.p1 | 21.153 | 190 | 1 | 13 | 5 | 13 | 5 | 47.9 | 17.4 | 323.31 | 176 | 33 | 31.6% | 6.8% |
| c6373_g1_i1.p1 | 119.64 | 1078 | 1 | 2 | 0 | 2 | 0 | 2.3 | 0 | 40.782 | 4 | 0 | 0.0% | 0.0% |
| c6458_g1_i1.p1 | 46.381 | 435 | 1 | 7 | 0 | 7 | 0 | 17.5 | 0 | 145.4 | 38 | 0 | 1.8% | 0.0% |
| c6656_g1_i1.p1 | 43.007 | 399 | 1 | 1 | 3 | 1 | 3 | 3.3 | 8.5 | 75.84 | 1 | 8 | 0.0% | 1.6% |
| c6797_g1_i1.p1 | 115.98 | 1048 | 1 | 0 | 2 | 0 | 2 | 0 | 1.7 | 10.995 | 0 | 2 | 0.0% | 0.0% |
| c6825_g1_i1.p3<br>c13559_g1_i1.p1 | 13.634 | 125 | 2 | 1 | 2 | 1 | 2 | 5.6 | 12.8 | 64.635 | 3 | 5 | 1.7% | 10.5% |
| c6945_g1_i1.p2 | 46.234 | 419 | 1 | 1 | 3 | 1 | 3 | 3.6 | 10.3 | 58.012 | 4 | 24 | 0.0% | 1.2% |
| c6963_g2_i1.p1 | 165.47 | 1543 | 1 | 4 | 2 | 4 | 2 | 2.1 | 1.6 | 55.502 | 9 | 11 | 0.1% | 0.2% |
| c7052_g1_i1.p1 | 24.213 | 227 | 1 | 16 | 7 | 2 | 4 | 45.4 | 31.7 | 323.31 | 147 | 27 | 32.4% | 7.3% |
| c7052_g1_i2.p1 | 23.965 | 227 | 1 | 15 | 5 | 1 | 2 | 40.1 | 25.1 | 140.21 | 15 | 9 | 4.1% | 2.0% |
| c7216_g1_i1.p1 | 25.446 | 231 | 1 | 3 | 0 | 3 | 0 | 16.9 | 0 | 19.32 | 7 | 0 | 0.4% | 0.0% |
| c8421_g1_i1.p1 | 59.681 | 570 | 1 | 4 | 1 | 4 | 1 | 10 | 2.3 | 323.31 | 62 | 2 | 1.6% | 0.0% |
| c926_g1_i1.p1 | 79.764 | 772 | 1 | 2 | 0 | 2 | 0 | 2.6 | 0 | 10.862 | 2 | 0 | 0.1% | 0.0% |

Table A.6: Summary of identifications and quantifications by semi-specific analysis (5% FDR) following parameter optimization following filtering of common contaminants (Stage 2).

| Protein ID | Low Peptide IDs | All high PEPs | Low protein score | Low sequence coverage | Low MS/MS count | N.D. in A | Highly enriched in B | Probable contaminant | Other | Filter |
|---|---|---|---|---|---|---|---|---|---|---|
| c10492_g1_i1.p1 | x | x | x | x | | x | x | | Only ID in tryp | x |
| c1275_g1_i1.p1 | x | x | x | x | | x | x | x | Only ID in semi | x |
| c1294_g1_i1.p1 | x | x | x | x | x | | | | Only ID in tryp | x |
| c1357_g1_i1.p1 | x | x | x | x | x | x | | | Only ID in tryp | x |
| c1505_g2_i1.p1 | | | | | | | | | | |
| c1613_g1_i1.p1 | x | (x) | x | (X) | | | | | | |
| c17161_g1_i1.p1 | x | (x) | x | (X) | x | x | | | Only ID in tryp | x |
| c17201_g1_i1.p1 | x | (x) | x | x | x | | | | Only ID in tryp | x |
| c17231_g1_i1.p1 | x | (x) | x | x | | | x | | | x |
| c17304_g1_i1.p1 | | | | | | | | | | |
| c17615_g1_i1.p1 | x | (x) | x | (x) | | | | | Only ID in tryp | |
| c231_g1_i1.p1 | x | (x) | x | | x | | | | Only ID in tryp | |
| c2364_g1_i1.p1 | | | | (x) | | | x | | | |
| c2556_g1_i1.p1 | | (x) | x | x | x | | | | Only ID in tryp | |
| c2788_g1_i1.p1 | x | (x) | | (x) | | x | x | | Only ID in semi | x |
| c3249_g1_i1.p1 | x | x | x | (x) | x | | | | Only ID in tryp | x |
| c3760_g1_i1.p1 | | | | (x) | | | | | Only ID in semi | |
| c4090_g1_i1.p1 | | (x) | x | | | | | | | |
| c4354_g1_i1.p1 | | | | | | | | | | |
| c4671_g1_i1.p2 | | (x) | | | | | | | | |
| c4757_g1_i1.p1 | x | | x | (x) | | x | x | | | x |
| c4921_g1_i1.p1 | | x | x | (x) | x | x | | | | x |
| c5168_g1_i1.p1 | x | | x | | x | x | | | | x |
| c5232_g1_i1.p1 | x | | | | | | | | | |
| c5952_g1_i1.p1 | x | x | x | (x) | | x | | | Only ID in semi | x |
| c6313_g1_i1.p1 | | | | | | | | | | |
| c6373_g1_i1.p1 | | | | x | | | | | | |
| c6458_g1_i1.p1 | | | | | | | | | | |
| c6656_g1_i1.p1 | | | | (x) | | | x | | | |
| c6797_g1_i1.p1 | x | (x) | x | x | x | x | | | | x |
| c6825_g1_i1.p3 | x | | | | | | x | x | | x |
| c6834_g1_i1.p3 | x | (x) | x | (x) | x | | | | Only ID in Tryp | |
| c6945_g1_i1.p2 | | | | (x) | | | x | x | | x |
| c6963_g2_i1.p1 | | (x) | | x | | | | | | |
| c7052_g1_i1.p1 | | | | | | | | | | |
| c7052_g1_i2.p1 | | | | | | | | | | |
| c7216_g1_i1.p1 | | | x | | | | | | | |
| c8389_g1_i1.p1 | x | (x) | x | x | x | | | | Only ID in Tryp | x |
| c8421_g1_i1.p1 | | | | | | | | | | |
| c926_g1_i1.p1 | | (x) | x | x | x | | | | | |

Table A.7: List of identified proteins using optimized analysis parameters (both samples and conditions included) and their adherence to the quality-based filtering criteria. Filtered proteins are marked in the right-most column.

| Condition | Total | Shared, all | Shared, sample | Tryptic, exclusive | Non-tryptic, exclusive |
|---|---|---|---|---|---|
| Tryptic, 1%FDR* | 219 | 48 (22%) | n.a | n.a | n.a. |
| Unspecific, 1%FDR | 279 | 21 (8%) | 100 (36%) | 119 (43%) | 60 (22%) |
| Unspecific, 5%FDR | 293 | 28 (10%) | 116 (40%) | 103 (35%) | 74 (25%) |
| Unspecific, 10%FDR | 328 | 31 (9%) | 134 (41%) | 85 (26%) | 109 (33%) |
| Semi-specific, 1%FDR | 285 | 26 (9%) | 135 (47%) | 84 (29%) | 66 (23%) |
| Semi-specific, 5%FDR | 312 | 39 (13%) | 168 (54%) | 51 (16%) | 93 (30%) |

Table A.8: Summary of identified peptides for the different search criteria used in the parameter optimization. The identified peptides are correlated between samples and between standard tryptic digestion (1% FDR) and the specified digestion criteria listed as found in the 4-way Venn diagrams (Figure A.3). Relative number of peptides (to the total number of identified peptides for the specified condition) is listed in parenthesis. *For tryptic digest at 1%FDR (standard settings), “shared, all” refers only to peptides shared between the two samples, and “n.a.” indicates that these metrics do not apply to 2-sample comparisons.

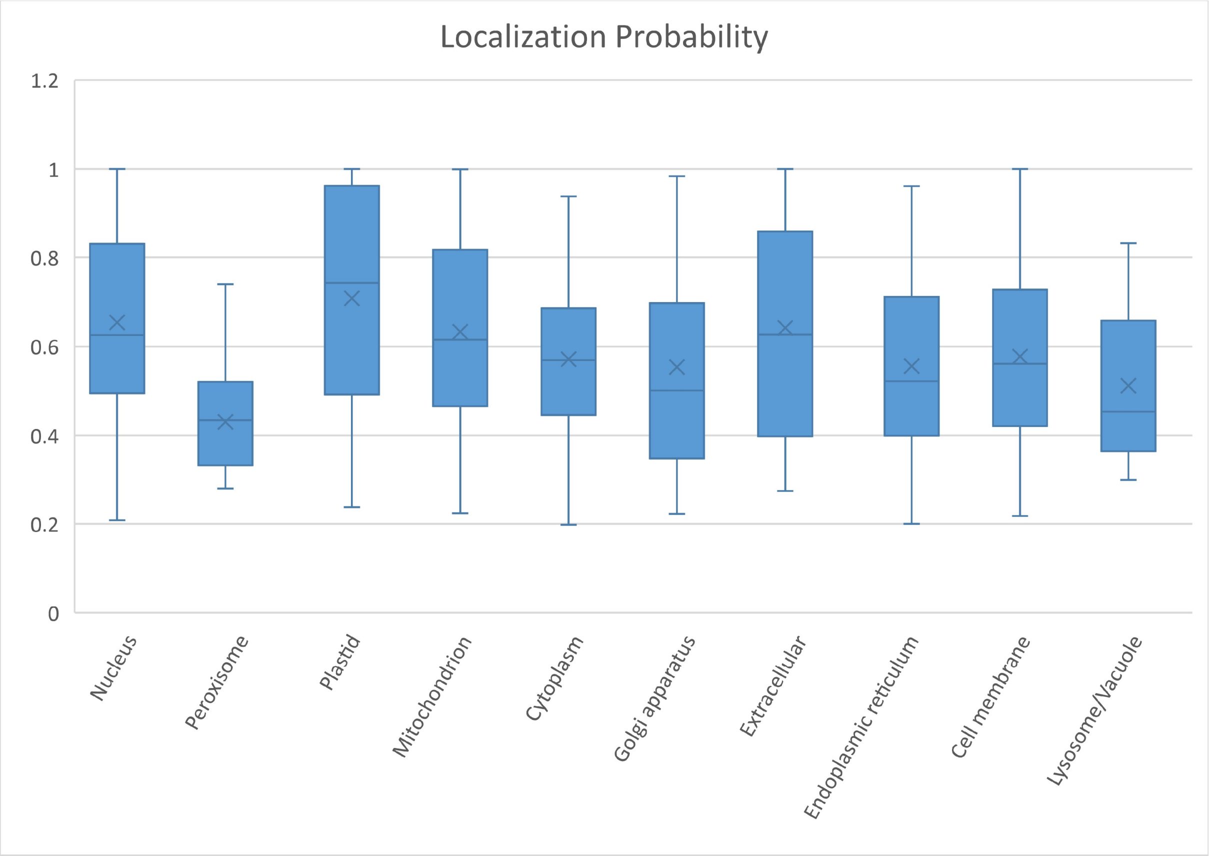


Figure A.1: Probability distribution for subcellular localization of the entire de-novo proteome for *E. denticulatum* using DeepLoc (Almagro Armenteros, Sønderby, Sønderby, Nielsen, & Winther, 2017)

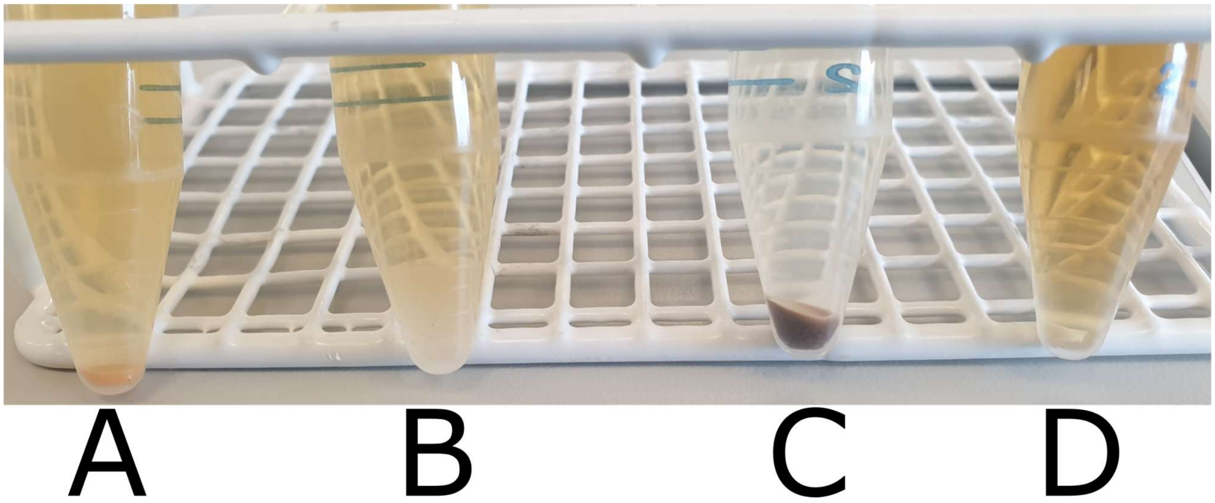


Figure A.2: Photograph of extract solutions and precipitate (after centrifugation) used for soluble protein quantification by Qubit. A: Extract A in $ddH_2O$. B: Extract A in 50mM $NH_4HCO_3$ with 0.2% SDS. C: Extract B in $ddH_2O$. D: Extract B in 50 mM $NH_4HCO_3$ with 0.2% SDS.  All solutions were made to a protein concentration of 2 mg/mL based on the supplied protein content by CP Kelco (Kjeldahl-N * 6.25).

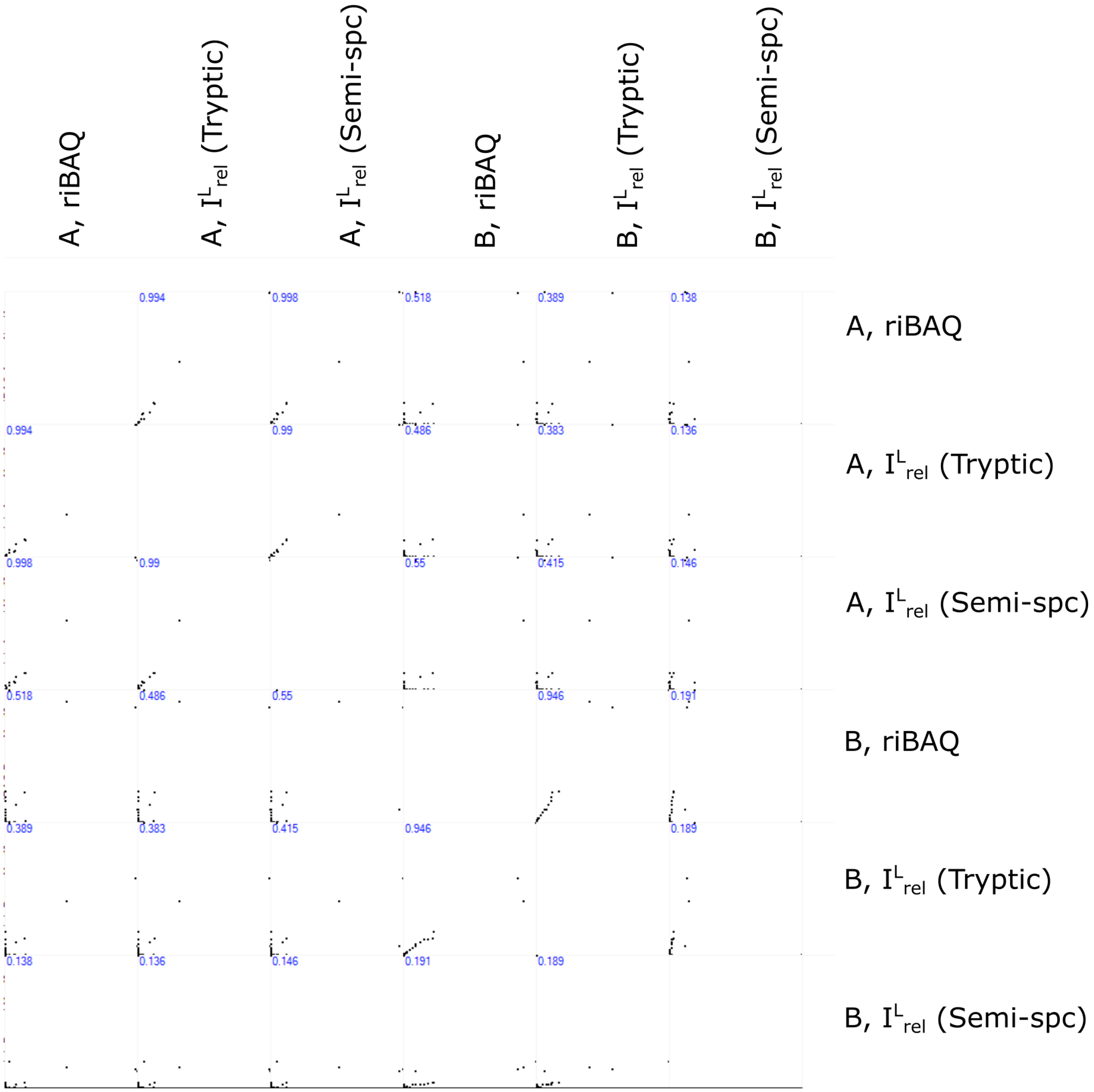


Figure A.3: Correlation of relative protein abundances between extracts (A and B), analysis conditions (tryptic and semi-specifc), and quantification method (riBAQ and $I^{L}_{rel}$) following filtering of common contaminants and re-quantification (Stage 2). Pearson Correlation Coefficients are shown in blue in the upper left corner of each sub-plot.

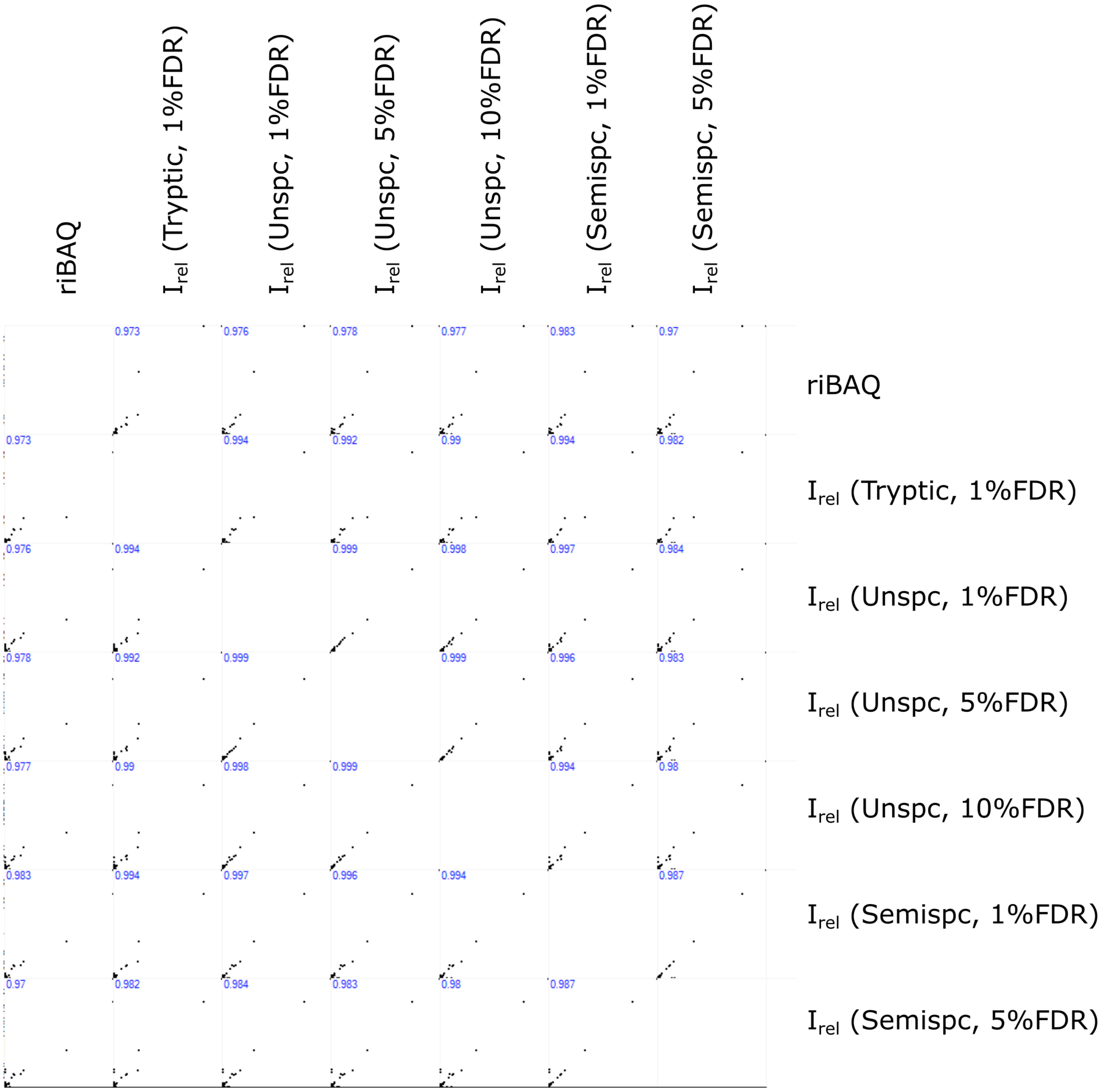


Figure A.4: Scatter plots of relative protein quantification for sample A using riBAQ from tryptic analysis and relative intensity for all searches. The Pearson Correlation Coefficient is indicated in blue at the upper left corner for each subplot.

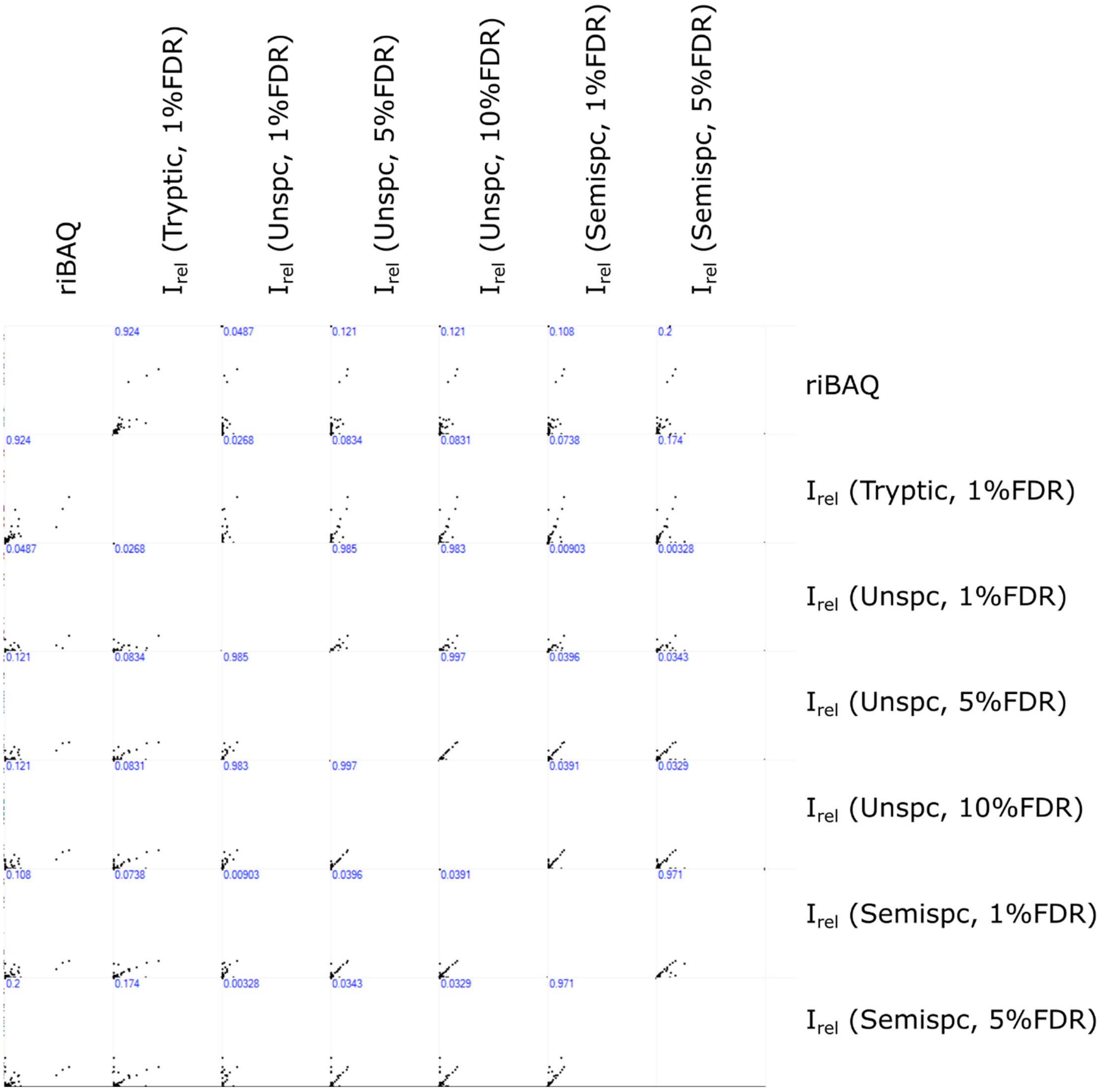


Figure A.5: Scatter plots of relative protein quantification for sample B using riBAQ from tryptic analysis and relative intensity for all searches. The Pearson Correlation Coefficient is indicated in blue at the upper left corner for each subplot

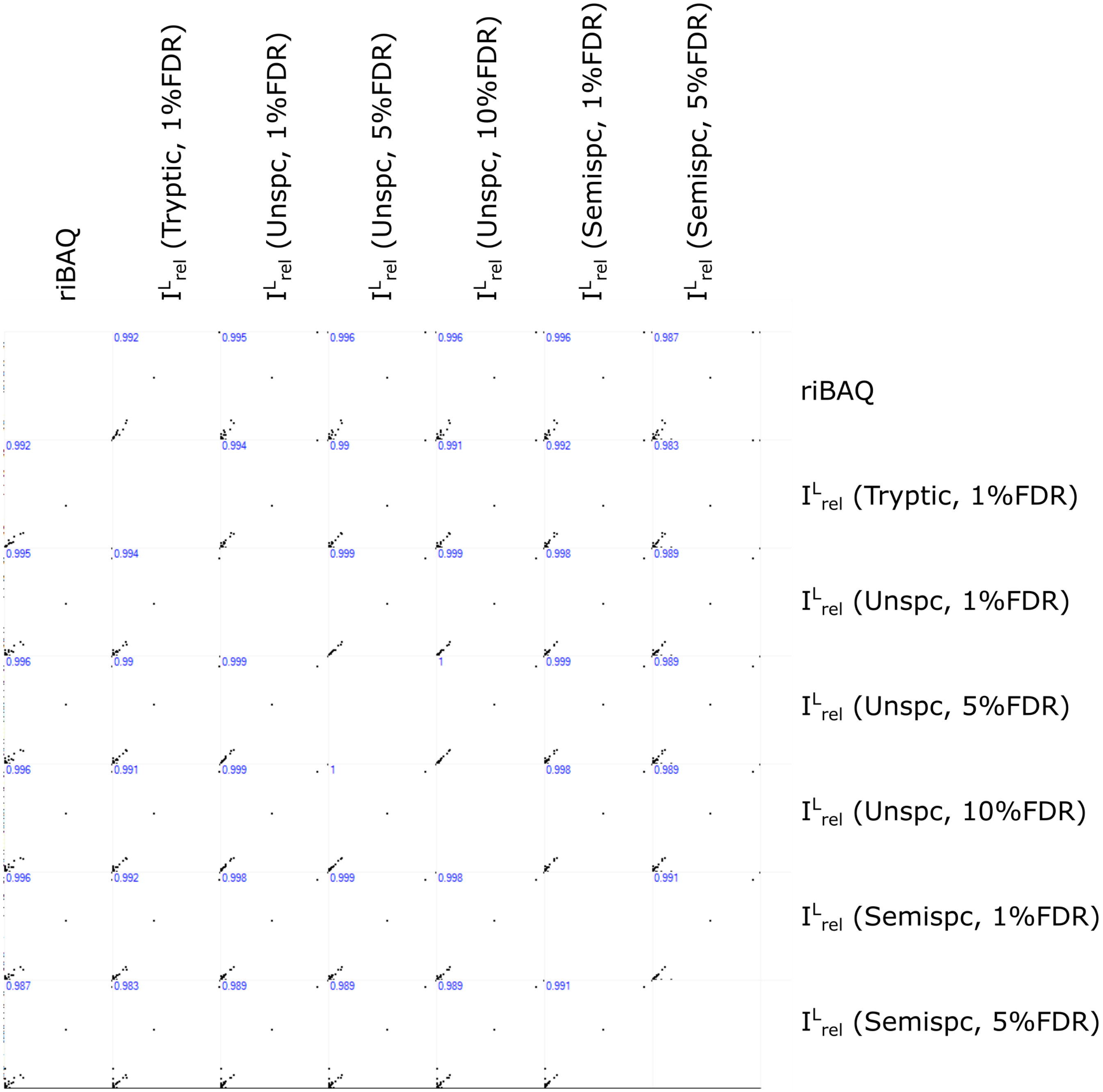


Figure A.6: Scatter plots of length-weighted relative protein quantification for sample A using riBAQ from tryptic analysis and length-weighted relative intensity for all searches. The Pearson Correlation Coefficient is indicated in blue at the upper left corner for each subplot

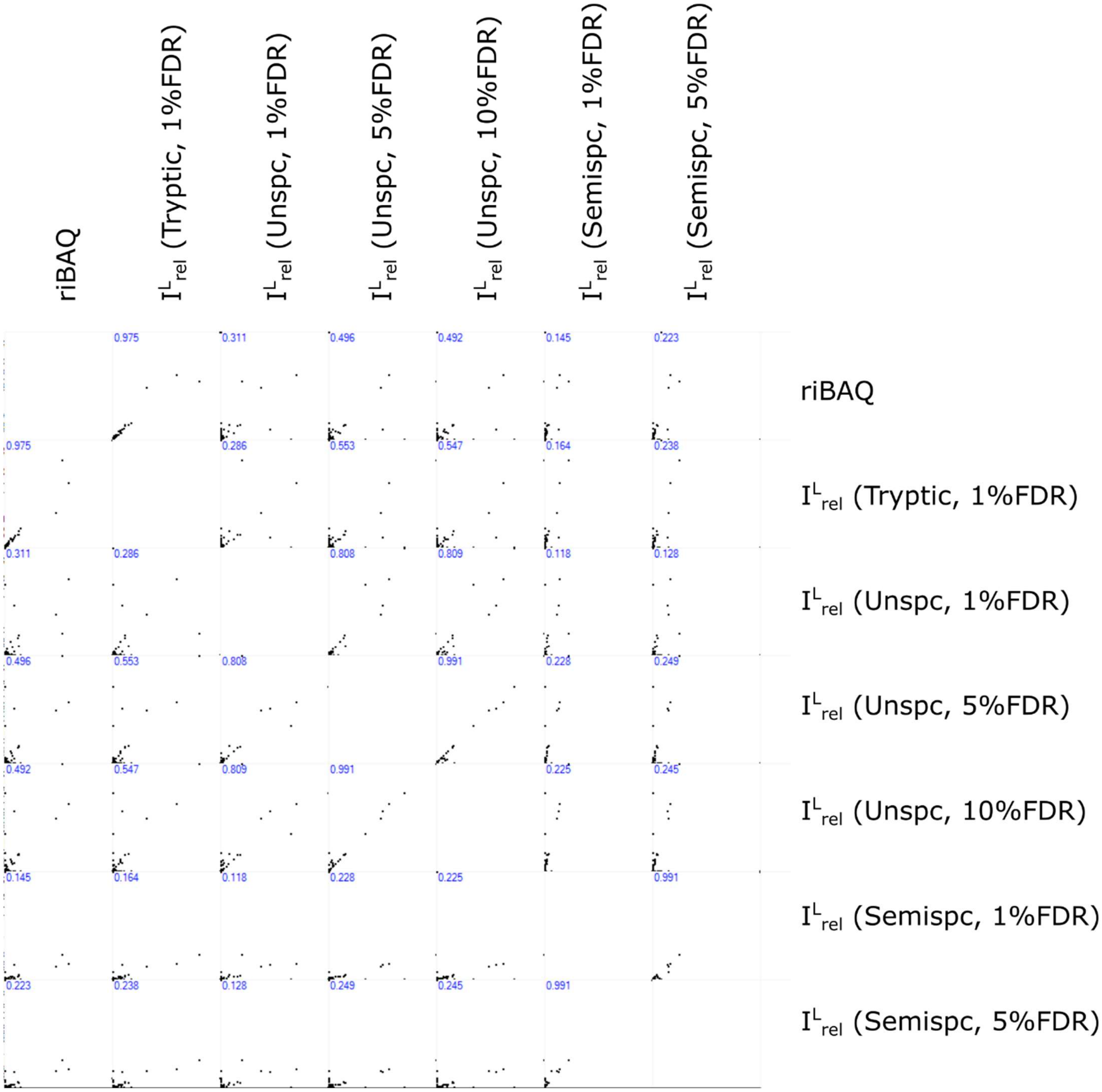


Figure A.7: Scatter plots of length-weighted relative protein quantification for sample B using riBAQ from tryptic analysis and length-weighted relative intensity for all searches. The Pearson Correlation Coefficient is indicated in blue at the upper left corner for each subplot

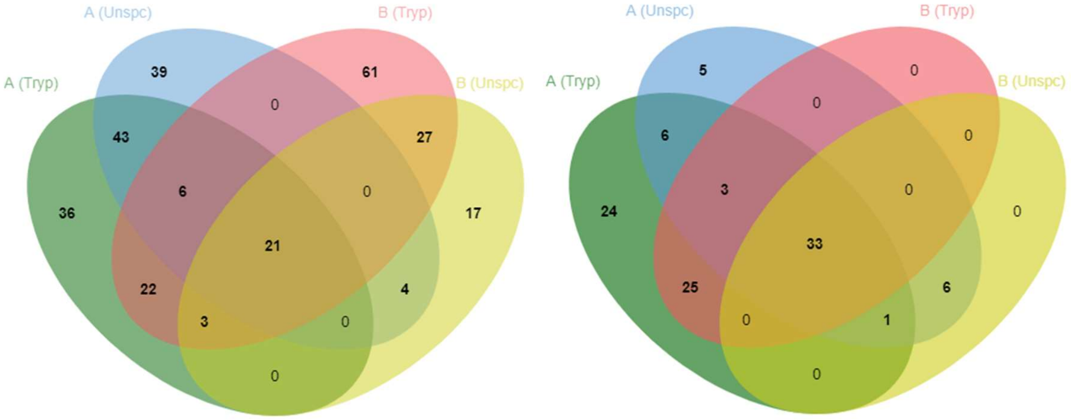


Figure A.8: Venn diagram of identified peptides (left) and proteins (right) in the A and B *E. denticulatum* protein extracts by LC-MS/MS using standard tryptic and unspecific digestion of the de-novo protein database in MaxQuant applying using 1% FDR on both peptide and protein level.

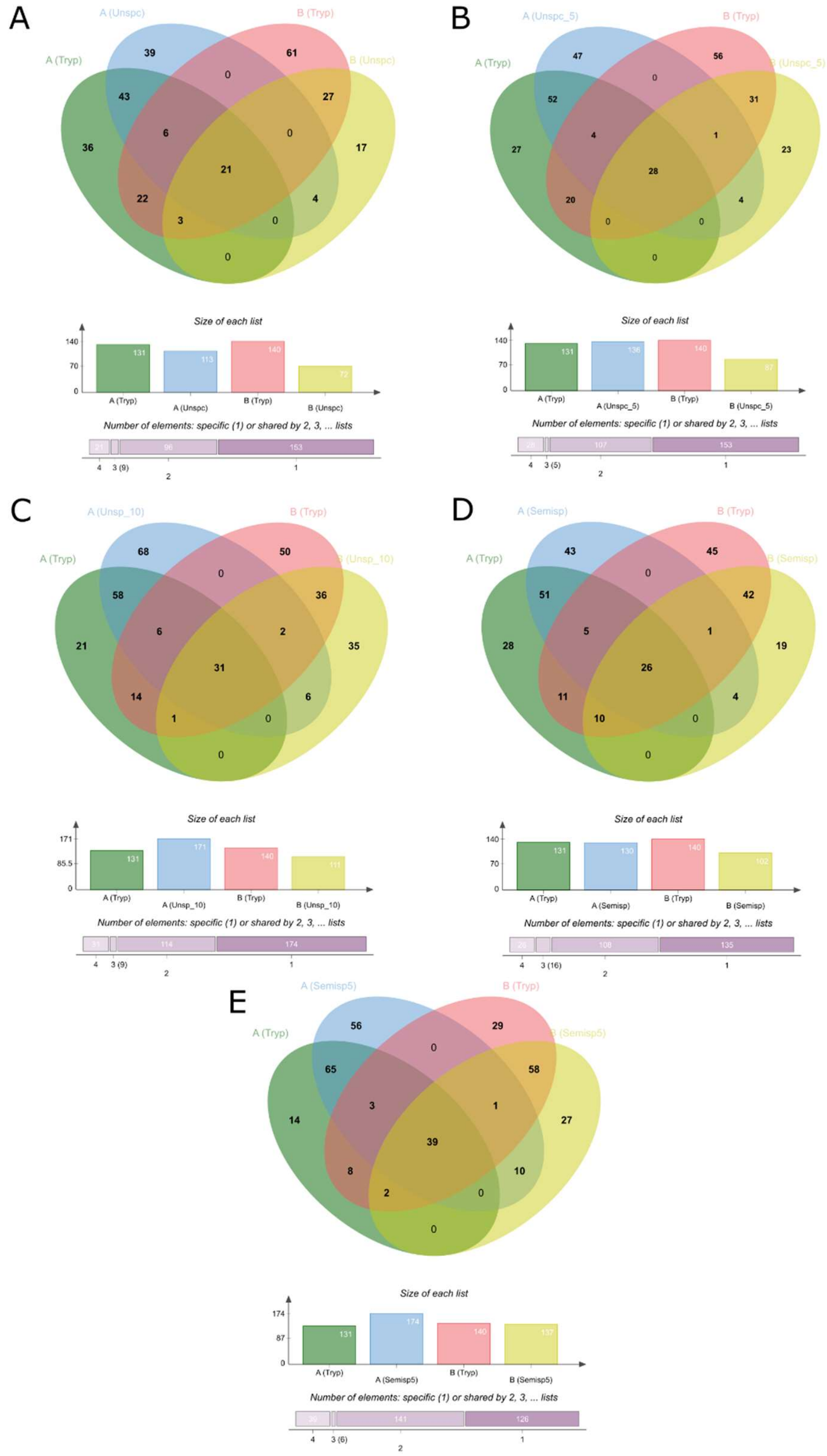


Figure A.9: 4-way Venn diagrams for peptides identified from the two extracts in MaxQuant. All Venn diagrams include identified protein by conventional tryptic analysis at 1% FDR combined with A) Unspecific, 1% FDR; B) Unspecific, 5% FDR; C) Unspecific, 10% FDR; D) Semi-specific, 1% FDR; E) Semi-specific, 5% FDR.

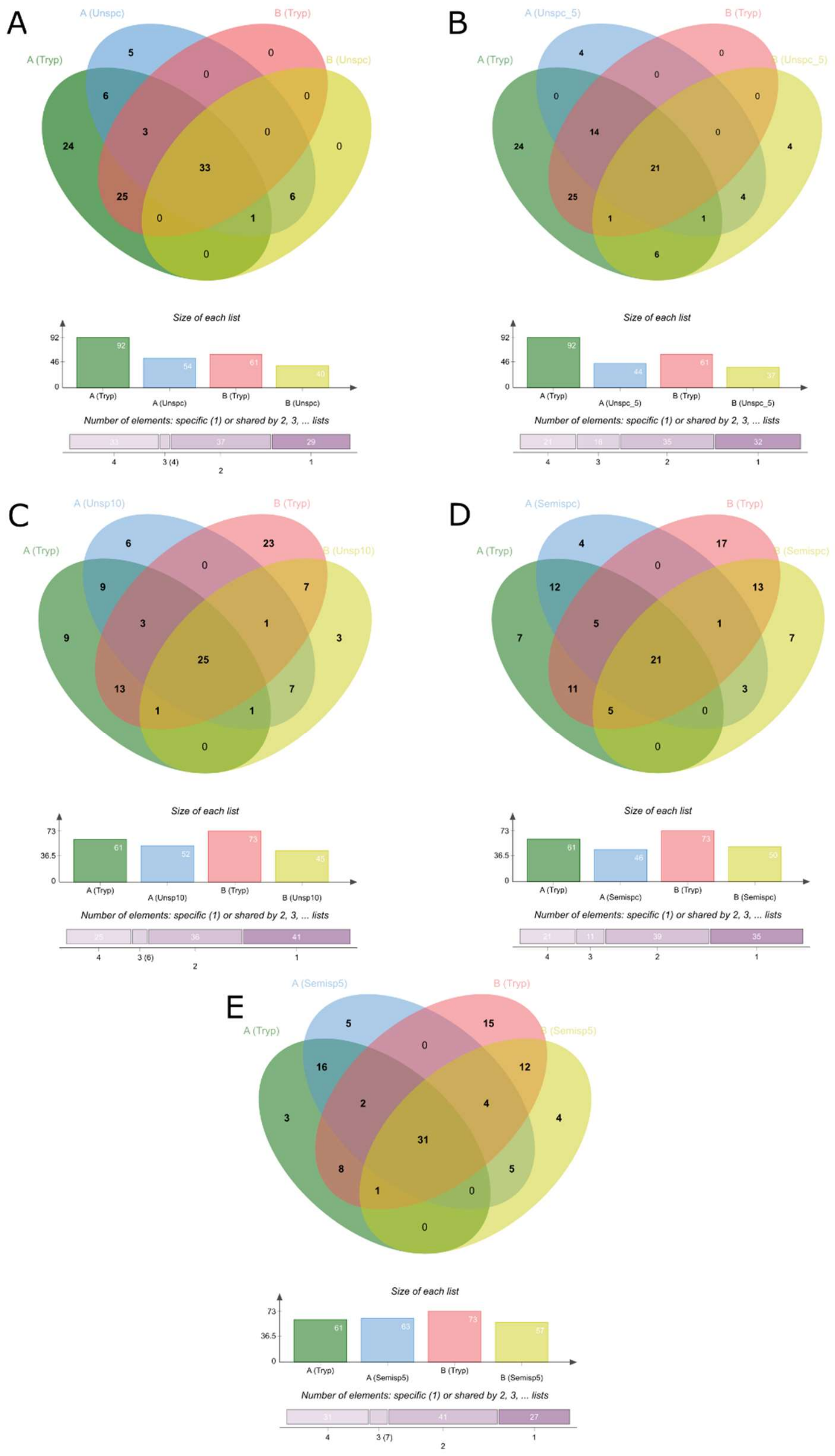


Figure A.10: 4-way Venn diagrams for proteins identified from the two extracts in MaxQuant. All Venn diagrams include identified protein by conventional tryptic analysis at 1% FDR combined with A) Unspecific, 1% FDR; B) Unspecific, 5% FDR; C) Unspecific, 10% FDR; D) Semi-specific, 1% FDR; E) Semi-specific, 5% FDR.

*References*


Almagro Armenteros, J. J., Sønderby, C. K., Sønderby, S. K., Nielsen, H., & Winther, O. (2017). DeepLoc: prediction of protein subcellular localization using deep learning. *Bioinformatics*, *33*(21), 3387–3395. https://doi.org/10.1093/bioinformatics/btx431

Bogdanow, B., Zauber, H., & Selbach, M. (2016). Systematic errors in peptide and protein identification and quantification by modified peptides. *Molecular and Cellular Proteomics*, *15*(8), 2791–2801. https://doi.org/10.1074/mcp.M115.055103

Chi, H., Liu, C., Yang, H., Zeng, W.-F., Wu, L., Zhou, W.-J., … He, S.-M. (2018). Open-pFind enables precise, comprehensive and rapid peptide identification in shotgun proteomics. *BioRxiv*, 285395. https://doi.org/10.1101/285395

Chi, H., Liu, C., Yang, H., Zeng, W. F., Wu, L., Zhou, W. J., … He, S. M. (2018). Comprehensive identification of peptides in tandem mass spectra using an efficient open search engine. *Nature Biotechnology*, *36*(11), 1059–1066. https://doi.org/10.1038/nbt.4236

Cox, Jurgen, Matic, I., Hilger, M., Nagaraj, N., Selbach, M., Olsen, J. V., & Mann, M. (2009). A practical guide to the maxquant computational platform for silac-based quantitative proteomics. *Nature Protocols*, *4*(5), 698–705. https://doi.org/10.1038/nprot.2009.36

Cox, Jürgen, Neuhauser, N., Michalski, A., Scheltema, R. A., Olsen, J. V., & Mann, M. (2011). Andromeda: A peptide search engine integrated into the MaxQuant environment. *Journal of Proteome Research*, *10*(4), 1794–1805. https://doi.org/10.1021/pr101065j

Fang, P., Liu, M., Xue, Y., Yao, J., Zhang, Y., Shen, H., & Yang, P. (2015). Controlling nonspecific trypsin cleavages in LC-MS/MS-based shotgun proteomics using optimized experimental conditions. *Analyst*, *140*(22), 7613–7621. https://doi.org/10.1039/c5an01505g

García-Moreno, P. J., Gregersen, S., Nedamani, E. R., Olsen, T. H., Marcatili, P., Overgaard, M. T., …

Jacobsen, C. (2020). Identification of emulsifier potato peptides by bioinformatics: application to omega-3 delivery emulsions and release from potato industry side streams. *Scientific Reports*, *10*(1), 690. https://doi.org/10.1038/s41598-019-57229-6

Gupta, N., & Pevzner, P. A. (2009). False discovery rates of protein identifications: A strike against the two-peptide rule. *Journal of Proteome Research*, *8*(9), 4173–4181. https://doi.org/10.1021/pr9004794

Jones, A. R., Siepen, J. A., Hubbard, S. J., & Paton, N. W. (2009). Improving sensitivity in proteome studies by analysis of false discovery rates for multiple search engines. *Proteomics*, *9*(5), 1220–1229. https://doi.org/10.1002/pmic.200800473

Lu, P., Vogel, C., Wang, R., Yao, X., & Marcotte, E. M. (2007). Absolute protein expression profiling estimates the relative contributions of transcriptional and translational regulation. *Nature Biotechnology*, *25*(1), 117–124. https://doi.org/10.1038/nbt1270

Mao, Y., Zhang, L., Kleinberg, A., Xia, Q., Daly, T. J., & Li, N. (2019). Fast protein sequencing of monoclonal antibody by real-time digestion on emitter during nanoelectrospray. *MAbs*, *11*(4), 767–778. https://doi.org/10.1080/19420862.2019.1599633

Swaney, D. L., Wenger, C. D., & Coon, J. J. (2010). Value of using multiple proteases for large-scale mass spectrometry-based proteomics. *Journal of Proteome Research*, *9*(3), 1323–1329. https://doi.org/10.1021/pr900863u

Tiwary, S., Levy, R., Gutenbrunner, P., Salinas Soto, F., Palaniappan, K. K., Deming, L., … Cox, J. (2019). High-quality MS/MS spectrum prediction for data-dependent and data-independent acquisition data analysis. *Nature Methods*, *16*(6), 519–525. https://doi.org/10.1038/s41592-019-0427-6

Tyanova, S., Temu, T., & Cox, J. (2016). The MaxQuant computational platform for mass spectrometry-based shotgun proteomics. *Nature Protocols*, *11*(12), 2301–2319. https://doi.org/10.1038/nprot.2016.136

Van Puyvelde, B., Willems, S., Gabriels, R., Daled, S., De Clerck, L., Vande Casteele, S., … Dhaenens, M. (2020). Removing the Hidden Data Dependency of DIA with Predicted Spectral Libraries. *PROTEOMICS*, *20*(3–4), 1900306. https://doi.org/10.1002/pmic.201900306

Wiśniewski, J. R. (2017). Label-Free and Standard-Free Absolute Quantitative Proteomics Using the “Total Protein” and “Proteomic Ruler” Approaches. In *Methods in Enzymology* (Vol. 585, pp. 49–60). https://doi.org/10.1016/bs.mie.2016.10.002

Wiśniewski, J. R., Ostasiewicz, P., Duś, K., Zielińska, D. F., Gnad, F., & Mann, M. (2012). Extensive quantitative remodeling of the proteome between normal colon tissue and adenocarcinoma. *Molecular Systems Biology*, *8*. https://doi.org/10.1038/msb.2012.44

Wiśniewski, J. R., Wegler, C., & Artursson, P. (2018). Multiple-Enzyme-Digestion Strategy Improves Accuracy and Sensitivity of Label- and Standard-Free Absolute Quantification to a Level That Is Achievable by Analysis with Stable Isotope-Labeled Standard Spiking. *Journal of Proteome Research*, *18*(1), acs.jproteome.8b00549. https://doi.org/10.1021/acs.jproteome.8b00549
